# Sensitive and modular amplicon sequencing of *Plasmodium falciparum* diversity and resistance for research and public health

**DOI:** 10.1101/2024.08.22.609145

**Authors:** Andrés Aranda-Díaz, Eric Neubauer Vickers, Kathryn Murie, Brian Palmer, Nicholas Hathaway, Inna Gerlovina, Simone Boene, Manuel Garcia-Ulloa, Pau Cisteró, Thomas Katairo, Francis Ddumba Semakuba, Bienvenu Nsengimaana, Hazel Gwarinda, Carla García-Fernández, William Louie, Endashaw Esayas, Clemente Da Silva, Debayan Datta, Shahiid Kiyaga, Innocent Wiringilimaana, Sindew Mekasha Fekele, Adam Bennett, Jennifer L. Smith, Endalamaw Gadisa, Jonathan B. Parr, Melissa Conrad, Jaishree Raman, Stephen Tukwasibwe, Isaac Ssewanyana, Eduard Rovira-Vallbona, Cristina M. Tato, Jessica Briggs, Alfredo Mayor, Bryan Greenhouse

## Abstract

**Background:** Targeted amplicon sequencing is a powerful and efficient tool for interrogating the *Plasmodium falciparum* genome, generating actionable data from infections to complement traditional malaria epidemiology. For maximum impact, genomic tools should be multi-purpose, robust, sensitive, and reproducible.

**Methods:** We developed, characterized, and implemented MAD^4^HatTeR, an amplicon sequencing panel based on Multiplex Amplicons for Drug, Diagnostic, Diversity, and Differentiation Haplotypes using Targeted Resequencing, along with a bioinformatic pipeline for data analysis. Additionally, we introduce an analytical approach to detect gene duplications and deletions from amplicon sequencing data. Laboratory control and field samples were used to demonstrate the panel’s high sensitivity and robustness.

**Results:** MAD^4^HatTeR targets 165 highly diverse loci, focusing on multiallelic microhaplotypes, key markers for drug and diagnostic resistance (including duplications and deletions), and *csp* and potential vaccine targets. The panel can also detect non-*falciparum Plasmodium* species. MAD^4^HatTeR successfully generated data from low-parasite-density dried blood spot and mosquito midgut samples, and detected minor alleles at within-sample allele frequencies as low as 1% with high specificity in high-parasite-density dried blood spot samples. Gene deletions and duplications were reliably detected in mono- and polyclonal controls. Data generated by MAD^4^HatTeR were highly reproducible across multiple laboratories.

**Conclusions:** The successful implementation of MAD^4^HatTeR in five laboratories, including three in malaria-endemic African countries, showcases its feasibility and reproducibility in diverse settings. MAD^4^HatTeR is thus a powerful tool for research and a robust resource for malaria public health surveillance and control.

## Background

Effective control and eventual elimination of *Plasmodium falciparum* malaria hinge on the availability and integration of data to inform research and public health strategies. Genomics can augment traditional epidemiological surveillance by providing detailed genetic information about infections^1^. Molecular markers of drug and diagnostic resistance can guide the selection of antimalarials and diagnostics, respectively^2–5^. Vaccine target sequences may shed light on vaccine efficacy and identify evidence of selective pressure^6^. Measures of genetic variation can provide insights into transmission intensity, rate and origin(s) of importation, and granular details of local transmission^7–14^. Differentiation of infections as either recrudescent or reinfections is critical for measuring outcomes of therapeutic efficacy studies that are used to guide antimalarial use worldwide^15–18^. Furthermore, the contribution of non-*falciparum* species to malaria burden is poorly characterized, and could complicate control and elimination efforts^19^.

To maximize public health and research utility, genomic methods should be robust and provide rich information from field samples, which may be low-density and are often polyclonal in malaria-endemic areas of sub-Saharan Africa^13,20–22^. While traditional genotyping methods of length polymorphisms and microsatellites can characterize malarial infections, they suffer from low sensitivity and specificity, and difficulties in protocol standardization^23–25^. Single nucleotide polymorphism (SNP) barcoding approaches have improved throughput, sensitivity and standardization^26,27^. However, the biallelic nature of most targeted SNPs limits their discriminatory power to compare polyclonal infections. Sequencing of short, highly variable regions within the genome containing multiple SNPs (microhaplotypes) provides multiallelic information that overcomes many of those limitations^28^. Microhaplotypes can be reconstructed from whole-genome sequencing (WGS) data or amplified by PCR and sequenced. Low abundance variants, especially in low-density samples, may be missed by WGS due to low depth of coverage. Amplicon sequencing offers much higher sensitivity and can target the most informative regions of the genome, increasing throughput and decreasing cost. Several Illumina-based multiplexed amplicon sequencing panels have been developed to genotype *P*. *falciparum* infections. SpotMalaria is a panel that genotypes 100 SNPs, most of which are biallelic, for drug resistance and diversity^26^. Pf AmpliSeq genotypes SNPs, currently focused on Peruvian genetic diversity, and also targets drug and diagnostic resistance markers^27^. Panels that target multiallelic microhaplotypes, including AMPLseq, provide greater resolution for evaluating polyclonal infections and also include drug resistance markers^29,30^. Nanopore-based amplicon panels enable the utilization of mobile sequencing platforms^31–33^. Thus, targeted amplicon sequencing is a flexible approach that has the potential to address multiple use cases. To fully realize this potential, a panel for research and public health would ideally include all necessary targets to answer a wide range of questions, while remaining modular to allow flexible allocation of sequencing resources.

Here, we developed MAD^4^HatTeR, an Illumina-compatible, multipurpose, modular tool based on Multiplex Amplicons for Drug, Diagnostic, Diversity, and Differentiation Haplotypes using Targeted Resequencing. MAD^4^HatTeR has 276 targets divided into two modules: A diversity module with 165 targets to assess genetic diversity and relatedness; and a resistance module consisting of 118 targets that cover 15 drug resistance-associated genes and assesses *hrp2*/*3* deletions, along with current and potential vaccine targets. The modules also include targets for non-*falciparum Plasmodium* species identification. We developed a bioinformatic pipeline to report allelic data, and implemented laboratory and bioinformatic methods in several sites, including countries in malaria-endemic sub-Saharan Africa. We then evaluated the panel’s performance on various sample types, including mosquito midguts, and showed that high quality data can be consistently reproduced across laboratories, including from polyclonal samples with low parasite density.

## Methods

### Participating laboratories

We generated data in five sites: the EPPIcenter at the University of California San Francisco (UCSF), in collaboration with the Chan Zuckerberg Biohub San Francisco, California; Infectious Diseases Research Collaboration (IDRC) at Central Public Health Laboratories (CPHL), Kampala, Uganda; Centro de Investigação em Saúde de Manhiça (CISM), Manhiça, Mozambique; National Institutes for Communicable Diseases (NICD), Johannesburg, South Africa; and Barcelona Institute for Global Health (ISGlobal), Barcelona, Spain. The procedures are described according to the workflows in San Francisco. Minor variations, depending on equipment availability, were implemented at other institutions.

### Amplicon panel design

We used available WGS data as of June 2021^3,30,34–42^ to identify regions with multiple SNPs within windows of 150-300 bp that lay between tandem repeats, using a local haplotype reconstruction tool (Pathweaver^43^). We compiled a list of drug resistance-associated and immunity-related SNPs (Tables 1 and 2) and identified regions of 150-300 bp between tandem repeats in and around *hrp2* and *hrp3* to assess diagnostic resistance-related deletions, as well as a region in chromosome 11 that is often duplicated in *hrp3*-deleted samples^44^. Paragon Genomics, Inc. designed amplification primers in multiplexed PCR using the Pf3D7 genome as a reference and related *Plasmodium* species and human genomes to design primers specific for *P*. *falciparum*. Genome verions and their GeneBank or RefSeq accessions for each species are: *P*. *falciparum* Pf3D7 (version=2020-09-01, GCA_000002765.3), *P*. *vivax* PvP01 (version=2018-02-28, GCA_900093555.2), *P*. *malariae* PmUG01 (version=2016-09-19, GCA_900090045.1), *P*. *ovale* PocGH01 (version=2017-03-06, GCA_900090035.2), *P*. *knowlesi* PKNH (version=2015-06-18, GCA_000006355.2) and *Homo sapiens* GRCh37 (GCA_000001405.14). In addition to the *P*. *falciparum* targets, we selected a target in the *ldh* gene (PF3D7_1325200) and its homologs in the other 4 *Plasmodium* species listed above for identification of concurrent infections with these species. To minimize PCR bias against longer amplicons, we restricted the design to amplicons of 225-275 bp, which can be covered with a significant overlap in paired-end sequencing in Illumina platforms with 300-cycle kits, except for targets around *hrp3* that needed to be 295-300 bp long to design primers successfully. We excluded or redesigned primers that contained more than 1 SNP (including non-biallelic SNPs) or indels in available WGS data or aligned to tandem repeats. To increase coverage of SNPs close to each other, we allowed for overlap in amplicons that targeted drug resistance and immunity-related markers. Primers were grouped in modules, as outlined in the results section (Figure 1 and Supplementary Table 1).

**Figure 1.**
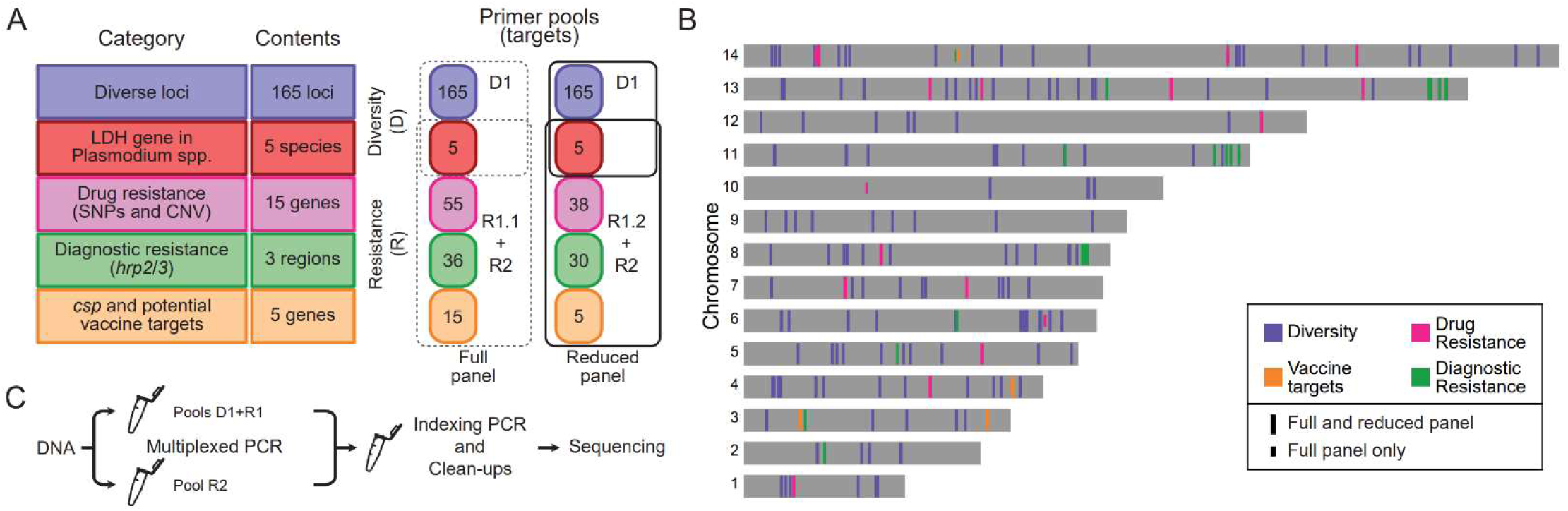
MAD^4^HatTer is a multi-purpose malaria amplicon sequencing panel. **A.** Primer pools to amplify targets in 5 categories are grouped into two modules (Diversity and Resistance). R1 refers to two primer pools: R1.1, the original pool, and R1.2, a reduced version of primer pool R1.1 designed to increase sensitivity. The recommended configuration to maximize information retrieval and sensitivity for low parasitemia samples are two mPCR reactions, one with D1 and R1.2 primers, and one with R2 primers (solid lines). Supplementary Tables 1-5 contain complete details on primer pools and targets. **B.** Chromosomal locations of all targets in the *P*. *falciparum* genome (not including non-*falciparum* targets). Note that the Diagnostic Resistance category includes targets in and around *hrp2* and *hrp3* as well as targets in chromosome 11 that are often duplicated when *hrp3* is deleted^44^ and length controls in other chromosomes. **C.** Simplified workflow for library preparation and sequencing, highlighting the need for two multiplexed PCR reactions when using primer pools R1 and R2 which are incompatible due to tiling over some genes of interest. A more detailed scheme can be found in Supplementary Figure 1, and a full protocol, including didactic materials, can be found online^78^.

**Table 1:** SNPs associated with antimalarial resistance. SNPs of interest used to optimize target primer design that are covered by primer pools R1.1, R1.2 or R2. The list excludes copy number variation markers, such as *plasmepsins* 2 and 3 (piperaquine) or *mdr1* (mefloquine). A full list of targets with the amino acid ranges they cover in each gene can be found in the Supplementary Table 4.

| Antimalarial<sup>†</sup> | Gene | Amino acids covered |
| --- | --- | --- |
| Chloroquine and piperaquine | *crt* | 72-76, 93, 97, 145, 218, 220, 271*, 326*<br>343, 350, 353, 356 |
| Chloroquine | *aat1* * | 135*, 162*, 185*, 230*, 238*, 380* |
| Piperaquine | *exo* | 415 |
| Chloroquine and lumefantrine | *mdr1* | 86, 184, 186, 371, 1034, 1042, 1246 |
| Sulfadoxine | *dhps* | 431, 436, 437, 540, 581, 613 |
| Pyrimethamine and proguanil | *dhfr* | 16, 51, 59, 108, 164 |
| Artemisinin | *kelch13* | 441, 446, 449, 458, 469, 476, 481, 493,<br>515, 527, 537, 538, 539, 543, 553, 561,<br>568, 574, 580, 622, 675 |
|  | *coronin* | 50, 100, 107 |
|  | *fd* | 193 |
|  | *arps10* | 127, 128 |
|  | *mdr2* | 463, 484, 515 |
|  | PF3D7_1322700 | 236 |
|  | *pib7* | 1484 |
|  | *ubp-1* | 1525 |
|  | *pph* * | 1157* |

<sup>†</sup> Antimalarial with validated or candidate markers<sup>79</sup>, or with SNPs identified in GWAS studies<sup>80</sup>.
\* Included only in R1.1 and not in R1.2.

**Table 2:**
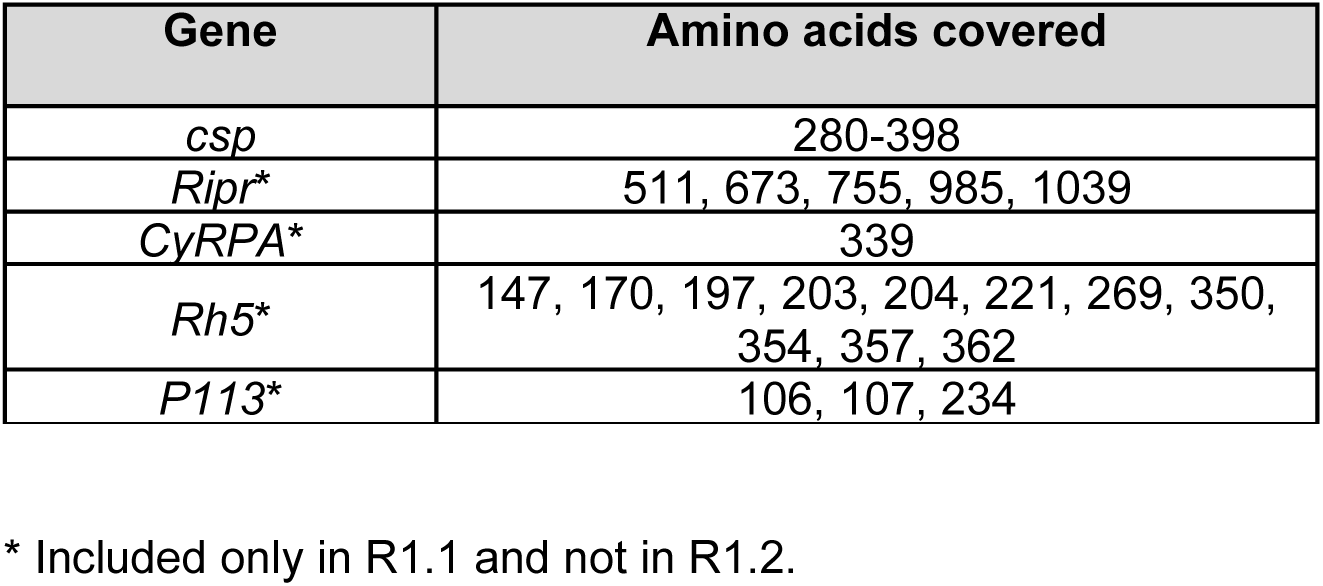
SNPs in *csp* and potential vaccine targets. SNPs of interest used to optimize target primer design that are covered by primer pools R1.1, R1.2 or R2.1. A full list of targets with the amino acid ranges they cover in each gene can be found in the Supplementary Table 4.

| Gene | Amino acids covered |
| --- | --- |
| *csp* | 280-398 |
| *Ripr* * | 511, 673, 755, 985, 1039 |
| *CyRPA* * | 339 |
| *Rh5* * | 147, 170, 197, 203, 204, 221, 269, 350, 354, 357, 362 |
| *P113* * | 106, 107, 234 |

\* Included only in R1.1 and not in R1.2.

### In silico panel performance calculations

Alleles were extracted from available WGS data as of July 2024^3,30,34–41,45^. SNPs, and microhaplotypes were reconstructed using Pathweaver^43^ for targets in MAD^4^HatTeR, SpotMalaria^26^, AMPLseq^30^, and AmpliSeq^27^. *In silico* heterozygosity was calculated using all allele calls in available WGS data. Principal coordinate analysis was performed on the binary distance matrix from presence/absence of alleles using alleles within loci present in both samples for each pair.

To assess statistical power of testing if two (potentially polyclonal) infections are related, we obtained within sample allele frequencies (WSAF) for the most variable SNP in each diversity target (165, 111 and 100 total SNPs for MAD^4^HatTeR, AMPLseq and SpotMalaria, respectively) or microhaplotypes (161, 128 and 135, respectively) from WGS data for each of the three panels, and simulated genotypes for mono- and polyclonal samples. In the simulations, complexity of infection (COI) were fixed and ranged from 1 to 5, and we included genotyping errors with a miss-and-split model^46^; missing and splitting parameters were 0.05 and 0.01, respectively. Between two samples, only a single pair of parasite strains was related with expected identity-by-descent (IBD) proportion varying from 1/16 to 1/2 (sibling level) to 1 (clones). We then analyzed these simulated datasets to obtain performance measures for combinations of a panel, COI, and a relatedness level: first, we estimated COI and allele frequencies using MOIRE^47^; we then used these to estimate pairwise interhost relatedness and test the hypothesis that two infections are unrelated at significance level of 0.05 with Dcifer^46^ and calculated power as the proportion of 1000 simulated pairs where the null hypothesis was correctly rejected.

### Samples

We prepared control dried blood spots (DBS) using *P*. *falciparum* laboratory strains. We synchronized monocultures in the ring stage. We made polyclonal controls by mixing cultured strains (3D7, Dd2 MRA-156 and MRA-1255, D6, W2, D10, U659, FCR3, V1/S, and HB3), all synchronized and ring-staged at various proportions. We mixed all monocultures and mixtures with uninfected human blood and serially diluted them in blood to obtain a range of parasite densities (0.1-100,000 parasites/μL). We spotted 20 μL of the mixture on filter papers and stored them at -20 ⁰C until processing.

Finger-prick DBS samples were collected in Northern Ethiopia between 2022 and 2023 as part of a mixed-methods study, which included a case-control study in two highland districts and cross-sectional surveys in one lowland district^48^. In the case-control study, samples were obtained from symptomatic patients presenting at health facilities. These included malaria cases - individuals who tested positive for *P*. *falciparum* and/or *P*. *vivax* using a rapid diagnostic test (Bioline Malaria Ag P.f/Pan by Abott, STANDARD Q Malaria P.f/P.v Ag by SD Biosensor, or First Response Malaria Ag. P.f./P.v. Card Test) – as well as test-negative controls who were later confirmed positive for malaria via qPCR. Cross-sectional surveys in lowland areas were conducted at agricultural worksites and in households in nearby villages. The DBS were air-dried, stored individually with desiccant, and kept at -20°C until laboratory processing. No patient data collected during sampling was used in this analysis. We analyzed DNA extracts from samples from previous studies, including 26 field samples from Ethiopia known to carry deletions in the *hrp2* and/or *hrp3* genes^3^, as well as 11 *P*. *falciparum* co-infections from Uganda containing *P*. *malariae* and *P*. *ovale*^49^. Finally, we analyzed publicly available data from 436 field samples from Mozambique^22^. The original works detail the sampling schemes and additional sample processing procedures. We used genomic DNA from *P*. *knowlesi* Strain H, obtained through BEI Resources, NIAID, NIH, contributed by Alan W. Thomas.

To assess performance of the assay for oocysts, we infected 9 *Anopheles gambiae* s.s. mosquitos via direct membrane feeding with blood taken from participants who were diagnosed with symptomatic malaria in transmission studies in Uganda. Briefly, mosquitos were fed via direct membrane feeding with blood taken from participants in a cohort study based in Nanongera and Busia districts (3 patients, 5 infected midguts) and from patients diagnosed with P. falciparum malaria at Masafu General Hospital in Busia district (4 patients, 4 infected midguts). ^50,51^ *P*. *falciparum* presence in blood samples was confirmed by varATS qPCR. Oocysts were detected and quantified using mercurochrome staining and microscopy as previously described^51^.

### Library preparation

We extracted DNA from control DBS and *P*. *falciparum*/*P*. *vivax* co-infections using the Chelex-Tween 20 method^52^. Mosquito midgut DNA was extracted from dissected midguts using the QIAGEN DNeasy blood and tissue DNA extraction kit as previously described^53^. *P*. *falciparum* parasite density was quantified in all samples, including midgut extracts, by varATS^54^ or 18S^55^ qPCR using standards made from DBS spotted with serial dilutions of cultured *P*. *falciparum* in uninfected blood (Supplementary Text). *P*. *vivax* was quantified by 18S qPCR as previously described^56^.

Libraries were made with a minor adaptation of Paragon Genomics’ CleanPlex Custom NGS Panel Protocol^57^ (Supplementary Text). A version of the protocol containing any updates can be found at https://eppicenter.ucsf.edu/resources. Library pools were sequenced in Illumina MiSeq, MiniSeq, NextSeq 550, or NextSeq 2000 instruments with 150 paired-end reads. We tested different amplification cycles and primer pool configurations. Based on sensitivity and reproducibility, the following are the experimental conditions we use as a default: primer pools D1+R1.2+R2; 15 multiplexed PCR cycles for moderate to high parasite density samples (equivalent to ≥ 100 parasites/μL in DBS) and 20 cycles for samples with lower parasite density; 0.25X and 0.12X primer pool concentration, respectively.

### Bioinformatic pipeline development and benchmarking

We developed a Nextflow-based^58^ bioinformatic pipeline to filter, demultiplex, and infer alleles from fastq files (Supplementary Text). Briefly, the pipeline uses cutadapt^59^ and DADA2^60^ to demultiplex reads on a per-amplicon basis and infer alleles, respectively. The pipeline further processes DADA2 outputs to mask low-complexity regions, generate allele read count tables, and extract alleles in SNPs of interest. We developed custom code in Python and R to filter out low-abundance alleles and calculate summary statistics from the data. The current pipeline version, with more information on implementation and usage, can be found at www.github.com/EPPIcenter/mad4hatter.

We processed the data presented in this paper with release 0.1.8 of the pipeline.

We evaluated pipeline performance by estimating sensitivity (ability to identify expected alleles) and precision (ability to identify only expected alleles) from monoclonal and mixed laboratory controls with different proportions of strains (Supplementary Text). We tested the impact of multiple parameters and features on allele calling accuracy, including DADA2’s stringency threshold OMEGA_A and sample pooling treatment for allele recovery, masking homopolymers and tandem repeats, and post-processing filtering of low abundance alleles. Masking removed false positives with the trade-off of masking real biological variation. We obtained the highest precision and sensitivity using sample pseudo-pooling, highly stringent OMEGA_A (10^−120^), and a moderate postprocessing filtering threshold (minor alleles of > 0.75%). These results indicate that bioinformatic processing of MAD^4^HatTeR data can be optimized to retrieve accurate sample composition with a detection limit of approximately 0.75% WSAF.

For analyses of allelic data from mixed controls, only samples with ≥ 90% of targets with > 50 reads (183 for diversity, and 165 for drug resistance markers) were included in the analysis. For drug resistance markers, only SNPs with variation between controls were included (20/91 codons from 12/22 targets). Within a sample, targets with less than 100 reads were excluded as alleles with a minor WSAF of 1% are very likely to be missed. The large majority of controls (122/183 and 162/165 for diversity and drug resistance markers, respectively) had very good coverage (at most 2 missing loci).

Species-specific *ldh* targets in the panel were used to identify non-*falciparum* species. Targets with less than 5 reads were filtered out. *P*. *ovale ldh* target sequences were extracted from the *P*. *ovale curtisi* (PocGH01, GCA_900090035.2) or *P*. *ovale wallikeri* (PowCR01, GCA_900090025.2) genomes using target primer sequences. Observed sequences were then aligned to these reference sequences using BLAST. Heterozygosity was estimated using MOIRE^47^ version 3.2.0.

### Deletions and duplications

We used the following laboratory strains to benchmark deletion and duplication detection using MAD^4^HatTeR data: *pfhrp2* deletions in Dd2 and D10, *pfmdr1* duplications in Dd2 and FCR3, *pfhrp3* deletion in HB3, and *pfhrp3* duplication in FCR3^44^. We also used a set of field samples from Ethiopia previously shown to have deletions in and around *pfhrp2* and *pfhrp3* at multiple genomic breakpoints^3^. For sensitivity analysis using field samples, we estimated COI using MOIRE^47^ and excluded polyclonal samples due to the uncertainty in their true genotypes. Two field samples were excluded from the analysis due to discordance in breakpoint classification, possibly due to sample mislabeling and sequencing depth, respectively.

We applied a generalized additive model (Supplementary Text) to account for target length amplification bias and differences in coverage across primer pools, likely due to pipetting error. We fit the model on controls known not to have deletions or duplications to obtain correction factors for targets of interest within sample batches. We then estimated read depth fold changes from data for each gene of interest (*pfhrp2*, *pfhrp3* and *pfmdr1*). We did not have sufficient data to validate duplications in plasmepsin 2 and 3.

For a subset of laboratory controls copy numbers were determined by qPCR using previously described methods for *pfmdr1*^61^, *pfhrp2*, and *pfhrp3*^62^.

## Results

### MAD^4^HatTeR is a multi-purpose tool that exploits P. falciparum genetic diversity

We designed primers to amplify 276 targets (Figure 1, Supplementary Tables 1-4) and separated them into two modules: (1) Diversity module, a primer pool (D1) targeting 165 high diversity targets and the *ldh* gene in *P*. *falciparum* and in 4 non-*falciparum Plasmodium* species (*P*. *vivax*, *P*. *malariae*, *P*. *ovale*, and *P*. *knowlesi*); and (2) Resistance module, comprised of two complementary and incompatible primer pools (R1 and R2) targeting 118 loci that genotype 15 drug resistance-associated genes (Table 1) along with *csp* and potential vaccine targets (Table 2), assess for *hrp2*/*3* deletion, and identify non-*falciparum* species. The protocol involves two initial multiplex PCR reactions, one with D1 and R1 primers, and another with R2 primers (Figure 1C, Supplementary Figure 1). After multiplexed PCR, subsequent reactions continue in a single tube.

Based on publicly available WGS data, *P*. *falciparum* targets in the diversity module, excluding *ldh*, had a median of 3 SNPs or indels (interquartile range [IQR] 2-5, N=165, Supplementary Table 5). Most (140/165) targets were microhaplotypes (containing > 1 SNP or indel). Global heterozygosity was high, with 35 targets with heterozygosity > 0.75 and 135 with heterozygosity > 0.5. Within African samples, heterozygosity was > 0.75 in 40 targets, > 0.5 in 132 targets, and we observed 2 to 20 unique alleles (median of 5, across a minimum of 3617 samples) in each target. MAD^4^HatTeR included more high-heterozygosity targets than other published panels (Figure 2A, Supplementary Figures 2 and 3). Additionally, MAD^4^HatTeR targets better resolved geographical structure globally, within Africa, and even within a country^63^ (Figure 2B).

**Figure 2.**
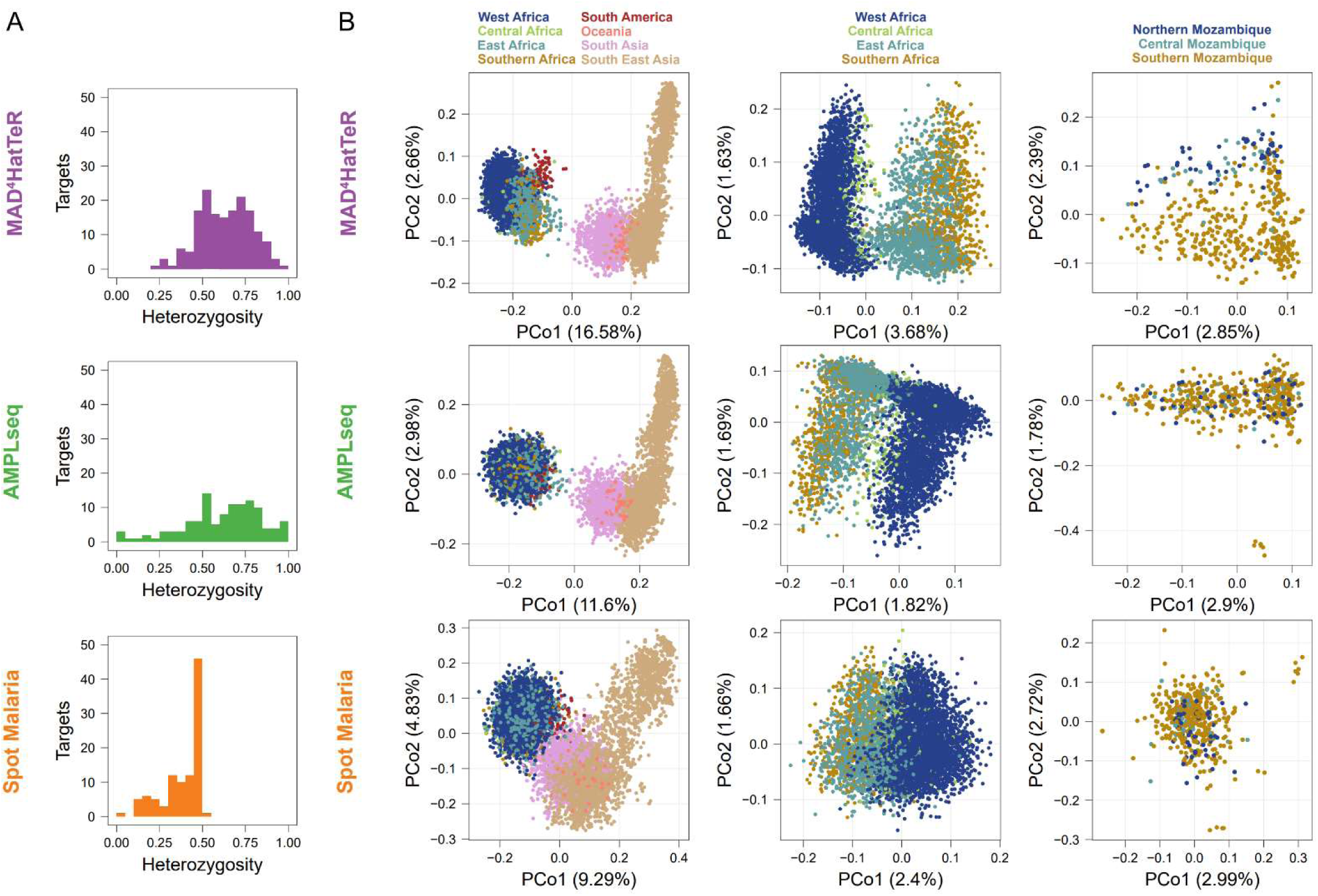
*In silico* analysis demonstrates that MAD^4^HatTer’s microhaplotypes capture high genetic diversity within African samples. We reconstructed alleles (microhaplotypes) from publicly available WGS data to estimate genetic diversity. For SpotMalaria, SNP barcodes are used instead of microhaplotypes based on intended design and current usage. We note that additional information may be present within the amplified targets if microhaplotype sequences are accurately identifiable with appropriate bioinformatic processing. As such, alternate results for microhaplotypes reconstructed for the targets that contain the SNPs in each of those two panels are shown in Supplementary Figure 2. **A.** Diversity module pool D1 includes more highly heterozygous targets than other published highly multiplexed panels. Only targets for diversity in each panel are included and heterozygosity is calculated for samples across Africa. **B.** We performed principal coordinate analysis on alleles on global, African or Mozambican data. The percentage of variance explained by each principal component is indicated in parentheses.

We next evaluated the power of the diversity module to detect interhost relatedness between parasites in pairs of simulated infections with COI ranging from 1 to 5. We selected one country from each of three continents with the most publicly available WGS data and used reconstructed genotypes for the analysis (Figure 3). MAD^4^HatTeR identified partially related parasites between polyclonal infections across a range of COI and geographic regions, and generally performed as well or better than the other panels evaluated. For example, in simulated Ghanaian infections sibling parasites (IBD proportion, r=½) were reliably detected with COI of 5 (82% power), half siblings (r=¼) in infections with COI of 3 (73% power), and less related parasites (r=⅛) were still identifiable with COI of 2 (53% power). When using independent SNPs instead of microhaplotypes, the power to identify related parasites between infections was much lower, irrespective of the panel. Constraining the panel to the 50 targets with the highest heterozygosity (mean heterozygosity of 0.8 ± 0.05) reduced the power to infer relatedness by as much as 50%, highlighting the value of highly multiplexed microhaplotype panels for statistical power.

**Figure 3.**
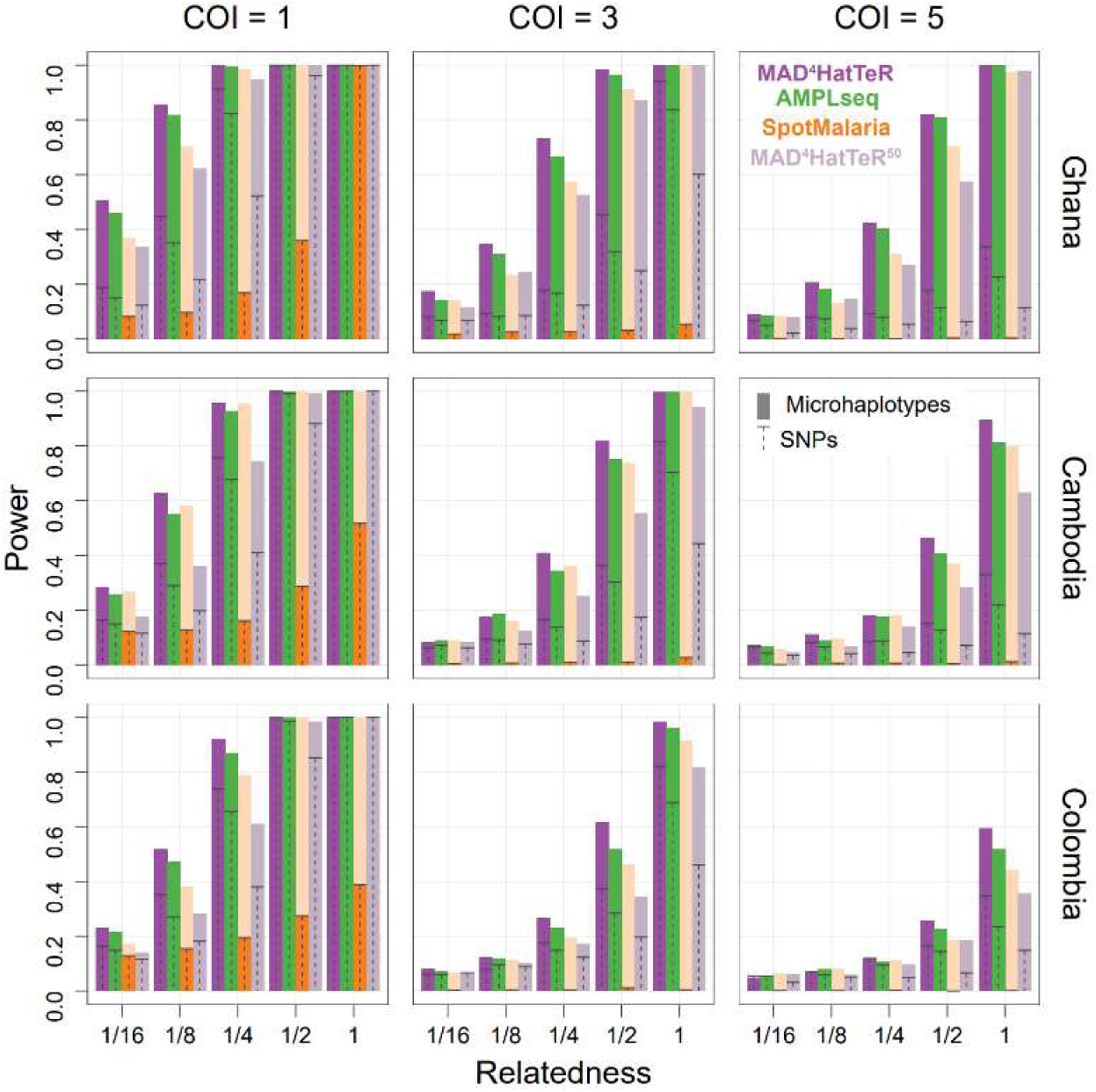
Power to identify relatedness of strains between infections is enhanced by highly multiplexed microhaplotypes. Simulated infections using population allele frequencies from available WGS data were used to estimate the power of testing if a pair of strains between infections is related. Countries in each of three continents with the most available WGS data were selected. Infections were simulated for a range of COI. Only one pair of strains between the infections was related with a given expected IBD proportion (r). The results were compared for reconstructed microhaplotypes and their most highly variable SNP for 3 panels (MAD^4^HatTeR, SpotMalaria and AMPLseq). Note that SpotMalaria bioinformatics pipeline outputs a 100 SNP barcode, and thus its actual power (dark orange) is not reflective of the potential power afforded by microhaplotypes (light orange). Additionally, the 50 most diverse microhaplotypes and their corresponding SNPs were used to evaluate the effect of down-sizing MAD^4^HatTeR (MAD^4^HatTeR^50^).

### MAD^4^HatTeR allows for genotyping of a variety of sample types and parasite densities

We evaluated MAD^4^HatTeR’s performance using dried blood spots (DBS) containing up to 7 different cultured laboratory strains each. Sequencing depth was lower for samples amplified with the original resistance R1 primer pool R1.1 than D1 (Supplementary Figure 4A), and primer dimers comprised 58-98% of the reads for R1.1 compared to only 0.1-4% for D1. We thus designed pool R1.2, a subset of targets from R1.1, by selecting the targets with priority public health applications and discarding the primers that accounted for a significant portion of primer dimers in generated data (Figure 1, Supplementary Table 2). Libraries prepared with pools containing R1.2 instead of R1.1 showed higher depth across the range of parasitemia evaluated (Supplementary Figure 4B). With the recommended set of primer pools (D1, R1.2, and R2), sequencing provided > 100 reads for most amplicons from DBS with > 10 parasites/µL, with depth of coverage increasing with higher parasite densities (Figure 4A). Samples with < 10 parasites/µL still yielded data albeit less reliably. Approximately 100,000 total unfiltered reads (the output of sample demultiplexing from a sequencing run) were sufficient to get good coverage across targets; on average, 95% of targets had at least 100 reads, and 98% had at least 10 reads (Supplementary Figure 4 C,D). While results indicate that the protocol provides consistently robust results, different experimental parameters may be optimal for different combinations of primer pools and sample concentration.

**Figure 4.**
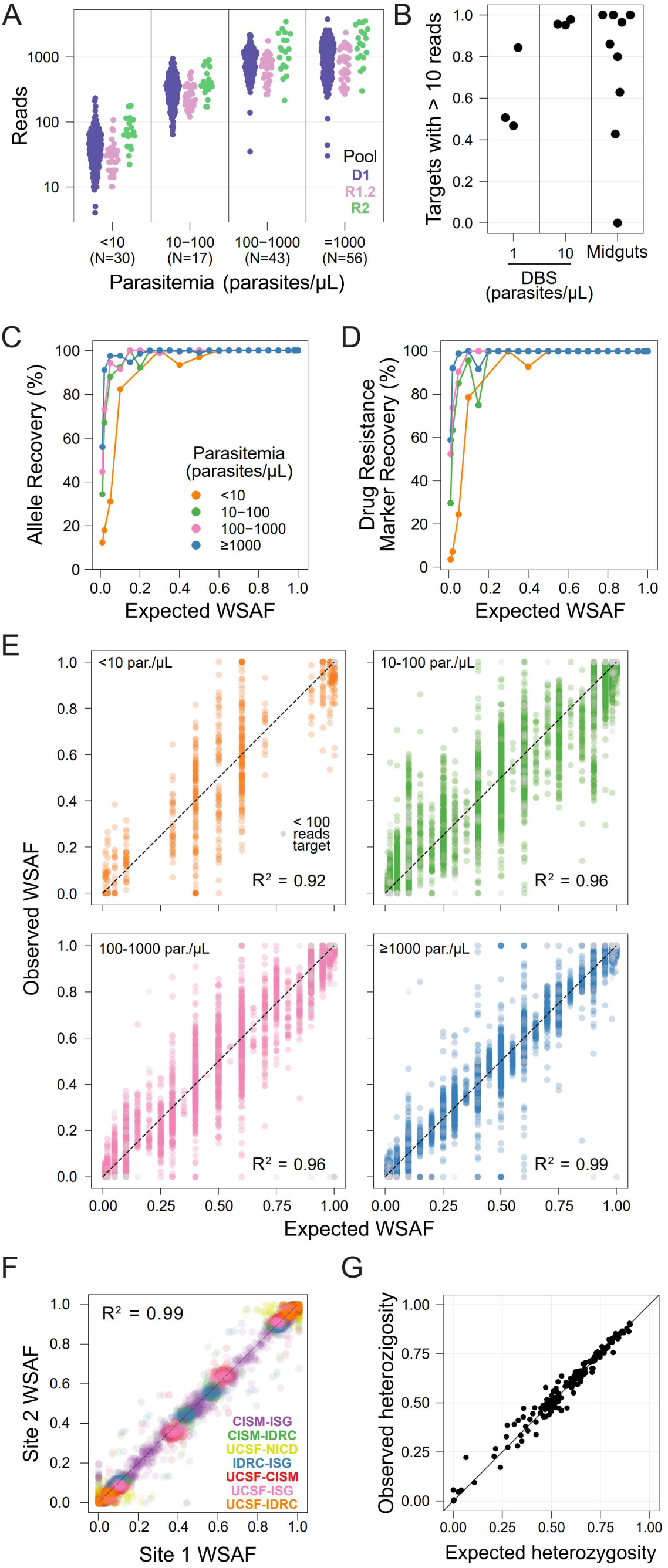
MAD^4^HatTeR produces reproducible and sensitive genetic data from a variety of samples. **A.** Mean read counts for each target in DBS controls (N in parenthesis in x-axis labels for each parasitemia). **B.** Proportion of targets with >10 reads in DBS controls with 1 and 10 parasites/μL and 9 midgut samples (median parasite density equivalent to 0.9 parasites/μL in a DBS). 10 targets that generally do not amplify well (>275 bp) were excluded. **C-D.** Recovery within-sample allele frequency (WSAF) in the diversity module for 161 loci across 183 samples (C), and biallelic SNPs in drug resistance markers across 20 codons in 165 samples (D). **E.** Observed WSAF in laboratory mixed controls of known expected WSAF. **F.** WSAF observed in libraries prepared and sequenced in different laboratories from the same DBS mixed control. Participating laboratories are the EPPIcenter at the University of California San Francisco (UCSF); Infectious Diseases Research Collaboration (IDRC), Uganda; Centro de Investigação em Saúde de Manhiça (CISM), Mozambique; National Institutes for Communicable Diseases (NICD), South Africa; and Barcelona Institute for Global Health (ISG), Spain. **G.** Observed heterozygosity in field samples from Mozambique^22^ and the respective expected heterozygosity for each target obtained from available WGS data (which does not include the MAD^4^HatTeR-sequenced field samples). False positives are excluded from C-G, as are targets with < 100 reads, except in E.

Depth of coverage per amplicon was highly correlated within technical replicates (Supplementary Figure 5A) with most deviations observed between primer pools. Importantly, coverage was also reproducible when the same samples were tested across five laboratories on 3 continents, with minor quantitative but negligent qualitative differences in coverage (Supplementary Figure 5B). Amplicon coverage was well balanced within a given sample, with differences in depth negatively associated with amplicon length (Supplementary Figure 6). Nine of the 15 worst-performing amplicons were particularly long (>297 bp, Supplementary Table 6). The other worst-performing amplicons covered drug resistance markers in *mdr1* and *crt* (neither covering *mdr1* N86Y or *crt* K76T), 2 high heterozygosity targets, and a target within *hrp2*. These results indicate that robust coverage of the vast majority of targets can be consistently obtained from different laboratories.

Given the high sensitivity of the method, we evaluated the ability of MAD^4^HatTeR to generate data from sample types where it is traditionally challenging to obtain high quality parasite sequence data. We amplified DNA extracted from nine infected mosquito midguts with a median *P*. *falciparum* DNA concentration equivalent to 0.9 parasites/µL from a DBS. On average, 58% of amplicons had ≥100 reads, 84% had ≥10 reads, and only one sample did not amplify (Figure 4B). These results are comparable to libraries from DBS controls with 1-10 parasites/µL from the same sequencing run, where 45-77% of amplicons with ≥100 reads. WSAF indicated that some of the mosquito midguts contained several genetically distinct *P*. *falciparum* clones. These data show the potential for applying MAD^4^HatTeR to study a variety of sample types containing *P. falciparum*.

### MAD^4^HatTeR reproducibly detects genetic diversity, including for minority alleles in low density, polyclonal samples

We used DBS controls containing 2 to 7 laboratory *P*. *falciparum* strains with minor WSAF ranging from 1 to 50% to evaluate sensitivity of detection and accuracy of WSAF estimation in the diversity pool D1. We optimized and benchmarked the bioinformatic pipeline to maximize sensitivity and precision, which included masking regions of low complexity (tandem repeats and homopolymers) to avoid capturing PCR and sequencing errors in allele calls. Sensitivity to detect minority alleles given that the locus amplified was very high, with alleles present at ≥ 2% reliably detected in samples with > 1,000 parasites/μL and at ≥ 5% in samples with > 10 parasites/μL (Figure 4C). For very low parasitemia samples (< 10 parasites/μL), sensitivity was still 82% for alleles expected at 10% or higher. Similar results were obtained for drug resistance markers targeted by pools R1.2 and R2 (Figure 4D). Overall precision (reflecting the absence of spurious alleles) was also high and could be increased by using a filtering threshold for minimum WSAF. Each sample had a median of 3 false positive alleles (mean = 4.4, N = 161 targets) above 0.75% WSAF, a median of 1 (mean = 2.5) false positives over 2%, and a median of 0 (mean = 0.7) over 5% (Supplementary Figure 7). A strong correlation between expected and observed WSAF was observed in the diversity module targets at all parasite densities and was stronger at higher parasite densities (R^2^=0.99 for > 1,000 parasites/µL Figure 4E).

Reproducibility is an important feature in generating useful data, particularly given differences in equipment and technique that often exists between laboratories. To evaluate this potential source of variation, we generated data for the same mixed-strain controls in five different laboratories on three continents. Reassuringly, the alleles obtained, along with their WSAF, were highly correlated (Figure 4F). Missed alleles in one or more laboratories were mostly present at < 2% within a sample. Finally, we tested MAD^4^HatTeR’s ability to recover expected diversity in field samples. Observed genetic heterozygosity in samples from Mozambique^22^ was correlated with expected heterozygosity based on available WGS data (Figure 4G, Supplementary Figure 8). These results highlight the reliability of MAD^4^HatTeR as a method to generate high quality genetic diversity data across laboratories.

### MAD^4^HatTeR provides data on copy number variations and detection of non-P. falciparum species

In addition to detecting sequence variation in *P. falciparum*, amplicon sequencing data can be used to detect gene deletions and duplications, as well as the presence of other *Plasmodium* species. We tested the ability of MAD^4^HatTeR to detect *hrp2* and *hrp3* deletions, and *mdr1* and *hrp3* duplications (laboratory strain FCR3 has a duplication in *hrp3*^44^) in DBS controls consisting of one or two laboratory strains, and field samples with previously known genotypes. We applied a generalized additive model to normalize read depth and estimate fold change across several targets per gene, accounting for amplicon length bias and pool imbalances, after using laboratory controls to account for batch effects, e.g. running the assay in different laboratories (Figure 5A, Supplementary Figure 9). The resulting depth fold changes for all loci assayed correlated with the expected sample composition (Figure 5B). At 95% specificity, sensitivity was 100% for all controls composed of > 95% strains with duplications or deletions (Figure 5C). Sensitivity was lower for samples with lower relative abundance of strains carrying duplications or deletions, although this could be increased with a tradeoff in specificity (e.g. if used as a screening test). Fold change data correlated well with quantification by qPCR, indicating that the data obtained from MAD^4^HatTeR are at a minimum semi-quantitative (Figure 5D). We could also correctly detect deletions in field samples from Ethiopia previously shown to be *hrp2*- or *hrp3*-deleted^3^, and correctly classify the genomic breakpoint profiles within the resolution offered by the targets included (Supplementary Figure 10). Finally, we detected *P*. *malariae* and *P*. *ovale* in 11 samples from Uganda known to contain the corresponding species, as previously determined by microscopy or nested PCR. We could distinguish *P*. *ovale wallikeri* from *P*. *ovale curtisi* based on the alleles in the target sequence. The assay’s sensitivity for detecting non-*falciparum* species was evaluated using a set of field samples from Ethiopia containing *P*. *falciparum* and *P*. *vivax*, with known parasite density for both species. Sensitivity depended on the *P*. *falciparum* to *P*. *vivax* ratio within the sample and was estimated at 96% for samples with more than 100 *P*. *vivax* 18S copies/μL (N = 148) and 90% for those with more than 10 18S copies/μL (N = 170) for samples with a *P*. *falciparum* to *P*. *vivax* ratio below 100 (Supplementary Figure 11). Furthermore, the *P*. *falciparum* to *P*. *vivax* ratio estimates obtained by qPCR and MAD^4^HatTeR were highly correlated. Specificity was 100% for all non-*falciparum* species, based on *P*. *falciparum* controls (N = 368). These data highlight the potential of MAD^4^HatTeR to capture non-SNP genetic variation and to characterize mixed species infections.

**Figure 5.**
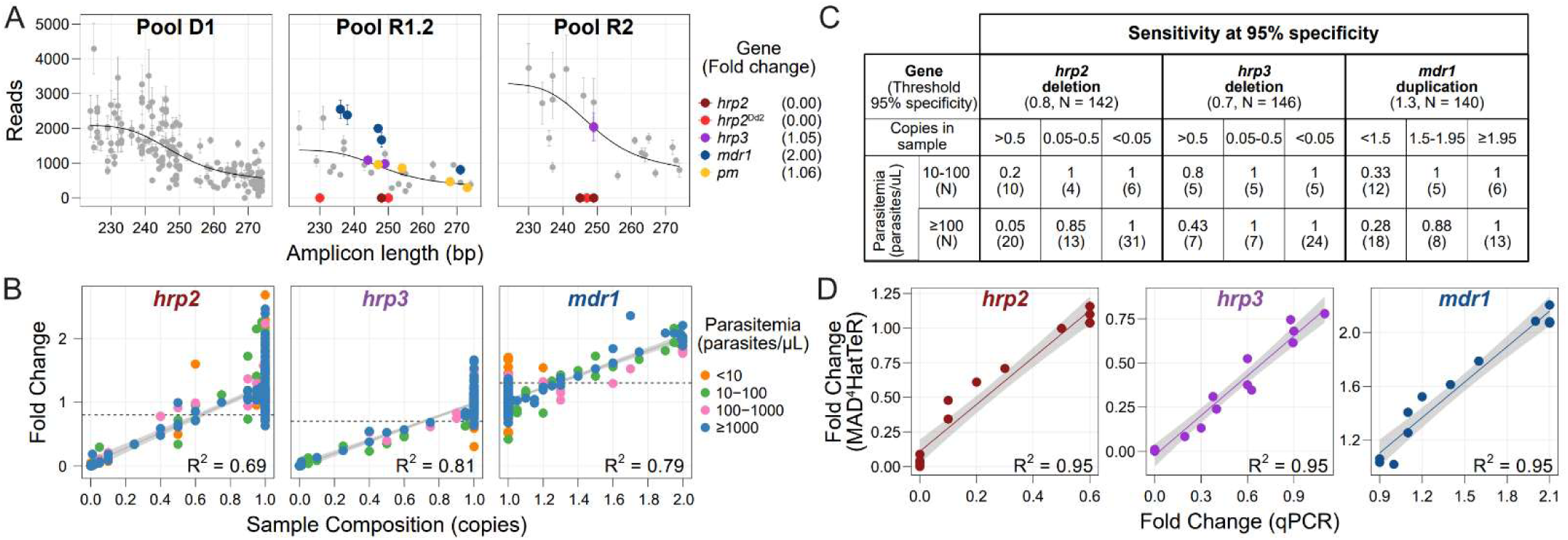
MAD^4^HatTeR can be used to screen for deletions and duplications. **A.** Technical replicates of Dd2 (a strain with *hrp2* deletion and *mdr1* duplication) with similar total reads were used to estimate fold changes in targets in and around *hrp2*, *hrp3*, *mdr1* and *plasmepsin2*/*3* (*pm*). A generalized additive model (black line) was applied to raw reads (Supplementary Figure 9) after correction by a control known not to have deletions or duplications in the genes of interest (3D7) to estimate fold changes in each of the genes. Note that there are two groups of *hrp2* targets, those that are deleted in field samples (*hrp2*) and those also deleted in Dd2 (*hrp2*^Dd2^). Mean reads and fold changes are shown (N = 3); error bars denote standard deviation. **B.** Estimated fold change for *hrp2*, *hrp3,* and *mdr1* loci in laboratory controls containing 1 or more strains at known proportions, or in field samples from Ethiopia^3^ with known *hrp2* and *hrp3* deletions. Sample composition is estimated as the effective number of copies present in the sample based on the relative proportion of the strain carrying a deletion or duplication. Fold changes are obtained using the targets highlighted in A. Fold changes for Dd2-specific targets are shown in Supplementary Figure 10. Linear regression and R^2^ values were calculated with data with parasitemia > 10 parasites/μL. The thresholds used to flag a sample as containing a duplication or deletion are shown in dashed black lines. **C.** Sensitivity in detecting *hrp2* and *hrp3* deletions and *mdr1* duplications in controls, and field samples from Ethiopia with known *hrp2* and *hrp3* deletions. Effective sample composition (copies in sample) is estimated as in B. Sensitivity was calculated using a threshold to classify samples with 95% specificity. Note that the small number of samples in the 0.05-0.5 copies range may be responsible for the paradoxical lower sensitivity for higher parasitemia samples. **D.** Estimated fold change for each gene correlates with qPCR quantification for the same samples.

## Discussion

In this study, we developed, characterized and deployed a robust and versatile method to generate sequence data for *P. falciparum* malaria genomic epidemiology, prioritizing information for public health decision-making. The modular MAD^4^HatTeR amplicon sequencing panel produces high-resolution data on genetic diversity, key markers for drug and diagnostic resistance, the C-terminal domain of the *csp* vaccine target, and presence of other *Plasmodium* species. MAD^4^HatTeR is highly sensitive, providing data for low parasite density DBS samples and detecting minor alleles at WSAF as low as 1% with good specificity in high parasite density samples; challenging sample types such as infected mosquitos were also successfully amplified. MAD^4^HatTeR has successfully generated data from field samples from Mozambique and Ethiopia, with particularly good recovery rates for samples with > 10 parasites/μL (∼90%)^22,64^. Deletions and duplications were reliably detected in mono- and polyclonal controls. The data generated by MAD^4^HatTeR are highly reproducible and have been reliably produced in multiple laboratories, including several in malaria-endemic countries. Thus, MAD^4^HatTeR is a valuable tool for malaria surveillance and research, offering policymakers and researchers an efficient means of generating useful data.

The 165 diversity and differentiation targets in MAD^4^HatTeR, of which the majority are microhaplotypes, can be used to accurately estimate within-host and population genetic diversity, and relatedness between infections. These data have promising applications: evaluating transmission patterns, e.g. to investigate outbreaks^3^; characterizing transmission intensity, e.g. to evaluate interventions^10,13,65^ or surveillance strategies^22^; classifying infections in low transmission areas as imported or local^11,66^; or classifying recurrent infections in antimalarial therapeutic efficacy studies as recrudescence or reinfections^18^. The high diversity captured by the current microhaplotypes could be further improved with updated WGS data to replace targets with relatively low diversity and amplification efficiency. Fully leveraging the information content of these diverse loci, which are particularly useful for evaluating polyclonal infections, requires bioinformatic pipelines able to accurately call microhaplotype alleles and downstream analysis methods able to incorporate these multi-allelic data. While some targeted sequencing methods and pipelines similarly produce microhaplotype data^30,32,67–69^, others only report individual SNPs, resulting in the loss of potentially informative data^26,27^ encoded in phased amplicon sequences. Many downstream analysis tools are similarly limited to evaluating data from binary SNPs^70–72^. Fortunately, methods to utilize these data are beginning to be developed, providing statistically grounded estimates of fundamental quantities such as population allele frequencies, COI^47^, and IBD^46^, and highlighting gains in accuracy and power provided by analysis of numerous highly diverse loci.

Multiple targeted sequencing tools designed with different use cases and geographies in mind are being used, raising questions about data compatibility. Comparing diversity metrics from data generated using different target sets is feasible, provided that the panels have equivalent performance characteristics and that the analysis methods appropriately account for differences such as allelic diversity^47^. Comparing genetic relatedness between infections evaluated with different panels, however, is limited to common loci. Over 25% of SNPs targeted by AMPLseq or SpotMalaria diversity targets were intentionally included in MAD^4^HatTeR. Other panels have less or no overlap^27,67,69^ (Supplementary Tables 9-10). Efforts to increase overlap between future versions of amplicon panels would facilitate more direct comparison of relatedness between infections genotyped by different panels.

MAD^4^HatTeR genotypes several key drug resistance markers as well as vaccine targets. The primer pool configuration recommended for optimized sensitivity covers markers of resistance to artemisinin, artemisinin-based combination therapy partner drugs, and other drugs used in treatment, chemoprevention, and other interventions. Additionally, it targets the C-terminal domain of *csp*, present in the RTS,S and R21 malaria vaccines currently recommended for use in children living in areas with moderate to high malaria transmission^73–75^. Other drug resistance markers and vaccine targets can be genotyped in high parasite density samples using the full primer pool configuration. Nevertheless, primer design and target prioritization have necessitated some exclusions. For example, the central repeat region of *csp*, also targeted by the RTS,S and R21 vaccines, is not covered. Future iterations of MAD^4^HatTeR should aim to include additional targets, such as evolving drug resistance markers and candidate vaccine targets.

Depth of coverage and amplification biases were reproducible across samples, with most deviations likely due pipetting volume differences and systematic differences in laboratory equipment and reagent batches. Detection of *hrp2*/*3* deletions and *mdr1* duplications was achieved by applying a model that accounts for these factors. MAD^4^HatTeR detected deletions and duplications in mono- and polyclonal samples, even at low parasitemia. Additional data and analytical developments could improve MAD^4^HatTeR’s performance in deletion and duplication analysis. The current approach does not make use of COI estimates for inference and relies on controls known not to have duplications or deletions in the target genes within each library preparation batch. While target retrieval was generally uniform, some samples showed target drop-off, indicating the need for multiple targets to avoid falsely calling a deletion. Nonetheless, in its current form, MAD^4^HatTeR serves as an efficient screening tool for identifying putative duplications and deletions, which can then be validated with gold-standard methodologies.

Continuous improvement of the allele-calling bioinformatic pipeline is planned to increase accuracy and usability. Masking of error-prone regions (e.g. homopolymers and tandem repeats) is useful in reducing common PCR and sequencing errors, but it also removes biological variation. This can be optimized by tailored masking of error hotspots, rather than uniformly masking all low-diversity sequences. To improve the detection of low-abundance alleles, we currently conduct a second inference round using alleles observed within a run as priors, but this approach may also increase the risk of incorporating low-level contaminant reads. Improvements in experimental strategies to detect and prevent cross-contamination^76^, along with post-processing filtering, could mitigate this. Additionally, curating an evolving allele database from ongoing empiric data generation could replace the run-dependent priors, thereby improving the accuracy and consistency of allele inference.

Integrating genomics into routine surveillance and developing genomic capacity in research and public health institutions in malaria-endemic countries is facilitated by efficient, cost-effective, reliable and accessible tools. MAD^4^HatTeR is based on a commercially available method for multiplexed amplicon sequencing^77^. As such, while primer sequences are publicly available (Supplementary Table 2), reagents are proprietary. However, procuring bundled, quality controlled reagents to generate libraries is straightforward, including for laboratories in malaria endemic settings. Procurement costs for laboratory supplies often vary significantly, making direct comparisons with other methods challenging, but we have found the method to be cost-effective compared with other methods. At the time of writing, the list price for all library preparation reagents, excluding plastics, consumables used for other steps (e.g. DNA extraction), sequencing costs, taxes, or handling, was $12-25 per reaction, depending on order volume. Sequencing costs can vary considerably based on the scale of sequencer used. For optimal throughput, we recommend multiplexing up to 96 samples using a MiSeq v2 kit to achieve results comparable to those shown here; much greater efficiency can be obtained with higher throughput sequencers.

This study includes data from five laboratories, three of which are located in sub-Saharan Africa. Beyond this study, MAD^4^HatTeR is also being used by four other African laboratories for applications ranging from estimating the prevalence of resistance-mediating mutations to characterizing transmission networks. Expertise and computational infrastructure for advanced bioinformatics and data analysis remains a challenge, with fewer users demonstrating autonomy in these areas compared to wet lab procedures. The robustness of the method, along with detailed training activities and materials (available online^78^), has facilitated easier implementation. Future developments could also expand accessibility, including adaptations for other sequencing platforms and panels targeting a smaller set of key loci for public health decision-making.

In summary, MAD^4^HatTeR is a powerful and fit-for-purpose addition to the malaria genomic epidemiology toolbox, well-suited for a wide range of surveillance and research applications.

## List of abbreviations

MAD^4^HatTeR: Multiplex Amplicons for Drug, Diagnostic, Diversity, and Differentiation Haplotypes using Targeted Resequencing
SNP: Single nucleotide polymorphism
WGS: Whole-genome sequencing
COI: Complexity of Infection
IBD: Identity-by-descent
DBS: Dried blood spot
WSAF: Within sample allele frequency

## Declarations

### Ethics approvals and consent to participate

Ethical approval for the study that generated the 26 field samples from Ethiopia^3^ was granted by the Ethiopia Public Health Institute (EPHI) Institutional Review Board (IRB; protocol EPHI-IRB-033-2017) and WHO Research Ethics Review Committee (protocol ERC.0003174 001).

Processing of de-identified samples and data at the University of North Carolina at Chapel Hill (UNC) was determined to constitute non-human subjects research by the UNC IRB (study 17-0155). The study was determined to be non-research by the Centers for Disease Control (CDC) and Prevention Human Subjects office (0900f3eb81bb60b9).

Study protocols for the study that generated the data for the 436 field samples from Mozambique^22^ were approved by the ethical committees of CISM and Hospital Clínic of Barcelona, and the Mozambican Ministry of Health National Bioethics Committee.

Ethical approval for the studies that included the collection of blood samples used in mosquito feeding assays was received from the Uganda Council of Science and Technology, Makerere University School of Medicine, the University of California, and the London School of Hygiene & Tropical Medicine.

Ethical approval for the study that collected *P*. *falciparum* and *P*. *vivax* samples in Northern Ethiopia in 2022-2023 was obtained from the National Research Ethical Review Committee, Addis Ababa, Ethiopia (reference number: 02/256/630/14), AHRI/ALERT Ethics Review Committee (protocol number: P0-08-22), Aklilu Lemma Institute of Pathobiology Institutional Research Ethics Review Committee (reference number: ALIPB IRERC/111/2015/23) and the WCG IRB approval (protocol number: 1769134-1; IRB tracking number: 20214694).

Participants, or guardians/parents in the case of minors, in all these studies provided written informed consent. All research was performed in accordance with relevant guidelines and regulations.

## Consent for publication

Not applicable.

## Availability of data and material

All data are available in the Sequencing Read Archive, accession code PRJNA1180199. Code is available in GitHub (https://github.com/EPPIcenter/mad4hatter and https://github.com/andres-ad/madh_utilities)

## Authors’ contributions

Designed the study: A.A.-D., N.H., A.B., J.L.S., E.G., B.G

Developed and benchmarked bioinformatic pipeline: A.A.-D., K.M., B.P., M.G.U., D.D.

Managed samples and data: A.A.-D, E.N.V, B.P., N.H, S.B, M.G.U., H.G., S.K, I.W., S.M.F, J.B.P., W.L., E.E.

Generated data: A.A.-D., E.N.V., S.B., P.C, T.K., F.D.S., B.N., H.G., C.G.F., C.D.S., A.E., S.M.F., W.L., E.E.

Analyzed data: A.A.-D., E.N.V., K.M., B.P., N.H., I.G., W.L.

Interpreted data: A.A.-D., E.N.V., K.M., B.P., N.H., I.G., M.G.U., M.C., J.R., S.T., I.S., E.R.-V., C.T., J.B., A.M., B.G., W.L

Drafted the manuscript: A.A.D., E.N.V., K.M., B.P., B.G

All authors reviewed the manuscript.

## Acknowledgments

We thank Phil Rosenthal and Amy Bei for their input in panel design. We also thank members of the EPPIcenter at UCSF, as well as the Rapid Response Team and the Genomics Platform at the Chan Zuckerberg Biohub for valuable discussions.

