## Supplementary Information for "Sensitive and modular amplicon sequencing of *Plasmodium falciparum* diversity and resistance for research and public health"

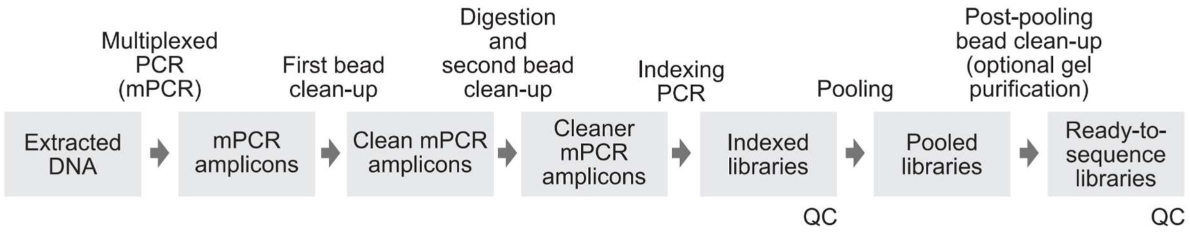


**Supplementary Figure 1.** Schematic of full protocol, which is based on Paragon Genomic's CleanPlex Amplicon Sequencing[1]. Suggested quality control (QC) steps by capillary electrophoresis are indicated. A full protocol, including didactic materials, can be found online[2].

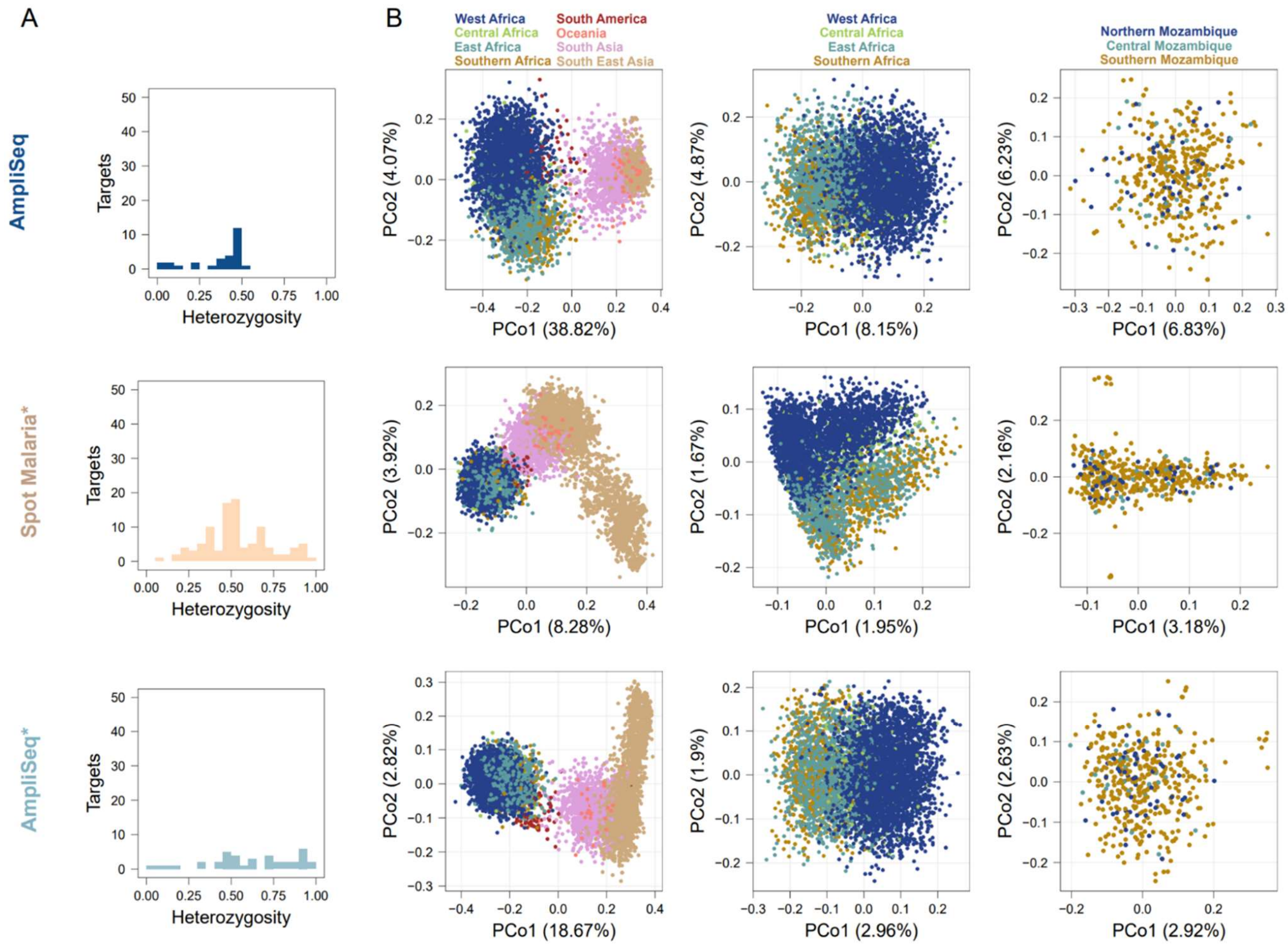


**Supplementary Figure 2**. Diversity module pool D1 includes more highly heterozygous targets than other published highly multiplexed panels.

**A**-**B**. SpotMalaria and AmpliSeq's use SNPs, not the full microhaplotypes. In its original publication, AmpliSeq used a 28 SNP barcode focused on Peruvian genetic diversity. Results for those 28 SNPs are shown in the first row. Microhaplotypes reconstructed from publicly available WGS data for the amplicons that contained the SNP barcodes in each of those panels (SpotMalaria* and AmpliSeq*) were used to calculate heterozygosity within Africa (**A**) and perform principal coordinate analysis (**B**).

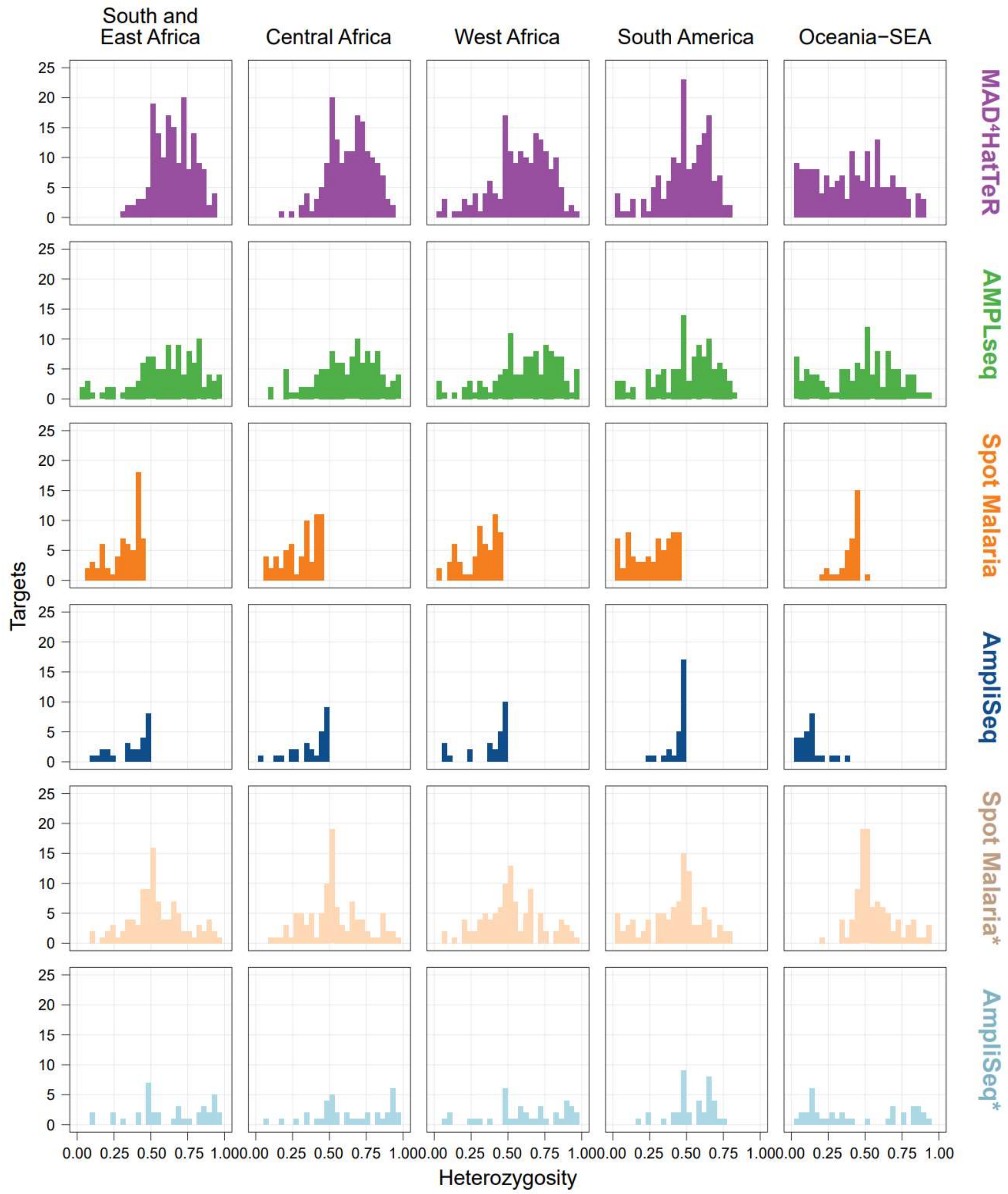


**Supplementary Figure 3**. Diversity module pool D1 includes more highly heterozygous targets than other published highly multiplexed panels across regions. Heterozygosity distributions for all panels in different regions. *SpotMalaria and AmpliSeq are shown both as SNP barcodes (intended and current use) and potential microhaplotypes (lighter color).

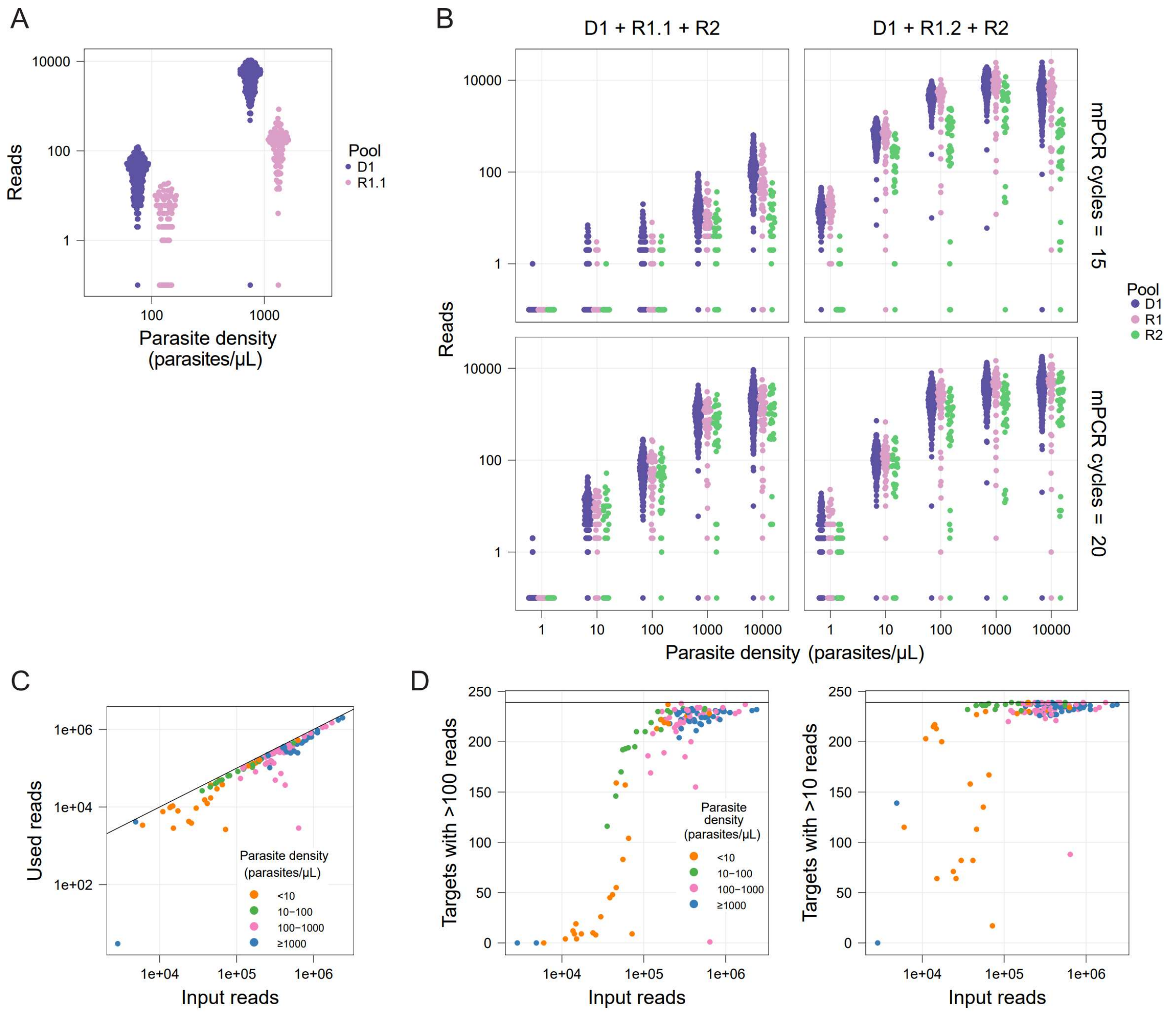


**Supplementary Figure 4**

**A**: Mean read count (N=2) for libraries prepared with either pool D1 or R1.1 show lower yields for R1.1, which was noted to result in more primer dimers.

**B**: Mean read count (N=2) for libraries prepared with either pools D1, R1.1 and R2, or D1, R1.2 (a subset of R1.1) and R2, show higher yields for libraries prepared with mixes including R1.2. Increasing multiplexed PCR cycles from 15 to 20 only increased yield for mixes containing R1.1

**C**: The majority of reads outputted by the sequencer (input reads) can be used for analysis as they pass all filters (for primer dimers, quality and alignment).

**D**: The number of targets with at least 100 (left) or 10 (right) reads passing all filters increases with the total number of reads outputted by the sequence (input reads). For samples with at least 100,000 reads, the number of targets with good coverage approximate the maximum number of targets in the sample (239, black horizontal line). Samples shown in C and D correspond to controls processed in different laboratories and sequenced in different runs, and samples with large (>10 fold) differences in coverage in the 2 reactions (likely due to pipetting error) were excluded. Note that many samples include controls with deletions in *hrp2*/*3* and thereby are not expected to reach 100% coverage. Additionally, longer amplicons and certain targets generally have lower coverage (Supplementary Figure 6).

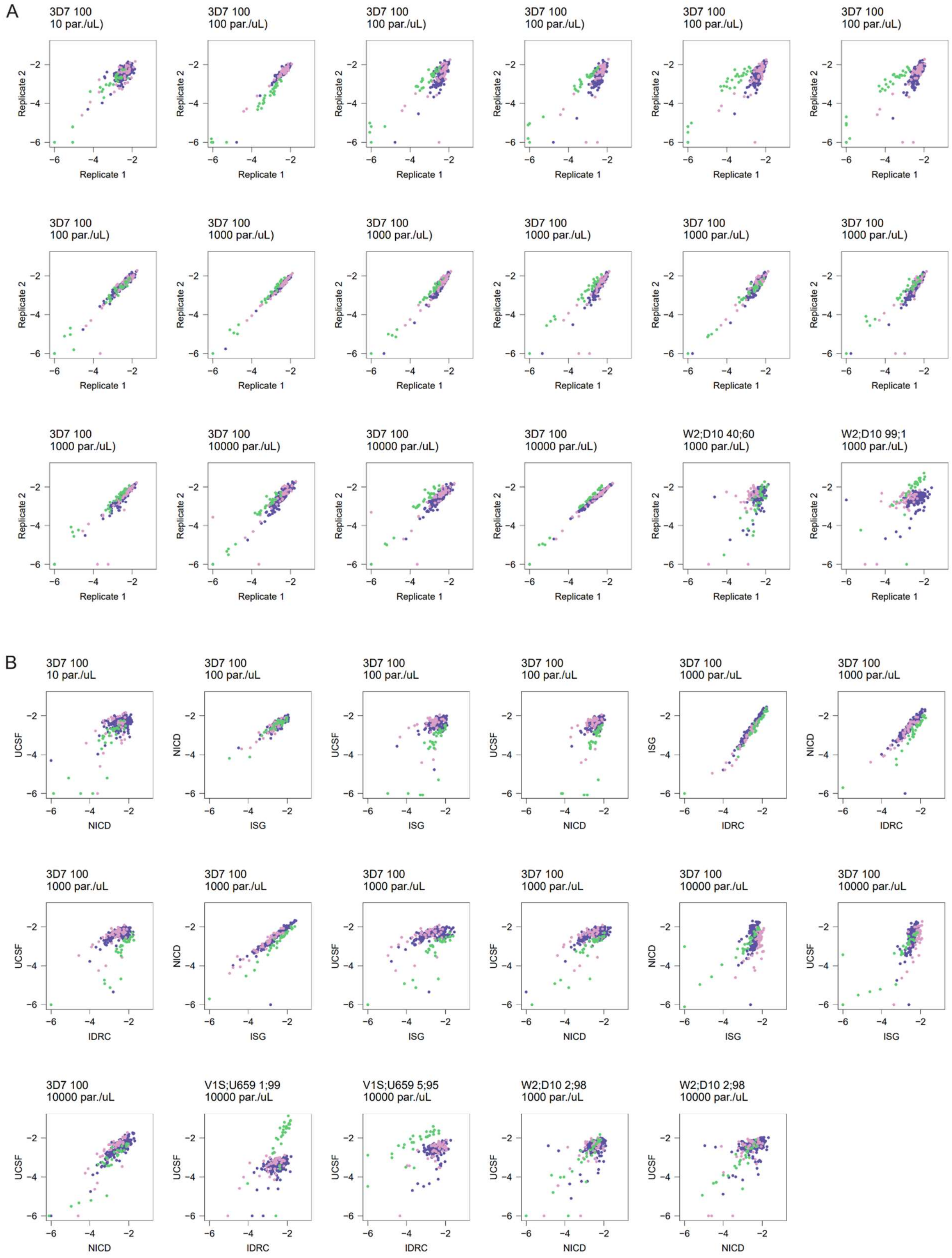
A
3D7 100
10 par./uL)
3D7 100
100 par./uL)
3D7 100
1000 par./uL)
3D7 100
10000 par./uL)
W2;D10 40;60
1000 par./uL)
W2;D10 99;1
1000 par./uL)
Replicate 1
Replicate 2
B
3D7 100
10 par./uL
3D7 100
100 par./uL
3D7 100
1000 par./uL
3D7 100
10000 par./uL
V1S;U659 1;99
10000 par./uL
V1S;U659 5;95
10000 par./uL
W2;D10 2;98
1000 par./uL
W2;D10 2;98
10000 par./uL
UCSF
NICD
ISG
IDRC

**Supplementary Figure 5**

Reproducibility for libraries prepared from the same DBS controls within the same laboratory or across laboratories. A subset of technical replicates prepared at the University of California San Francisco (UCSF) (**A**) or prepared in at least 2 different laboratories (**B**) are shown (ISGlobal: Barcelona Institute for Global Health, Spain; NICD: National Institute for Communicable Diseases, South Africa; IDRC: Infectious Diseases Research Collaboration at Central Public Health Laboratories, Uganda). Occasional systematic differences between coverage of pools likely represent pipetting error. Log10(reads) for each amplicon are shown in both axes. The DBS control strains are indicated with their composition (e.g. W2;D10 2;98 is 2% W2, 98% D10) and parasite density in parasites/µL (par./µL).

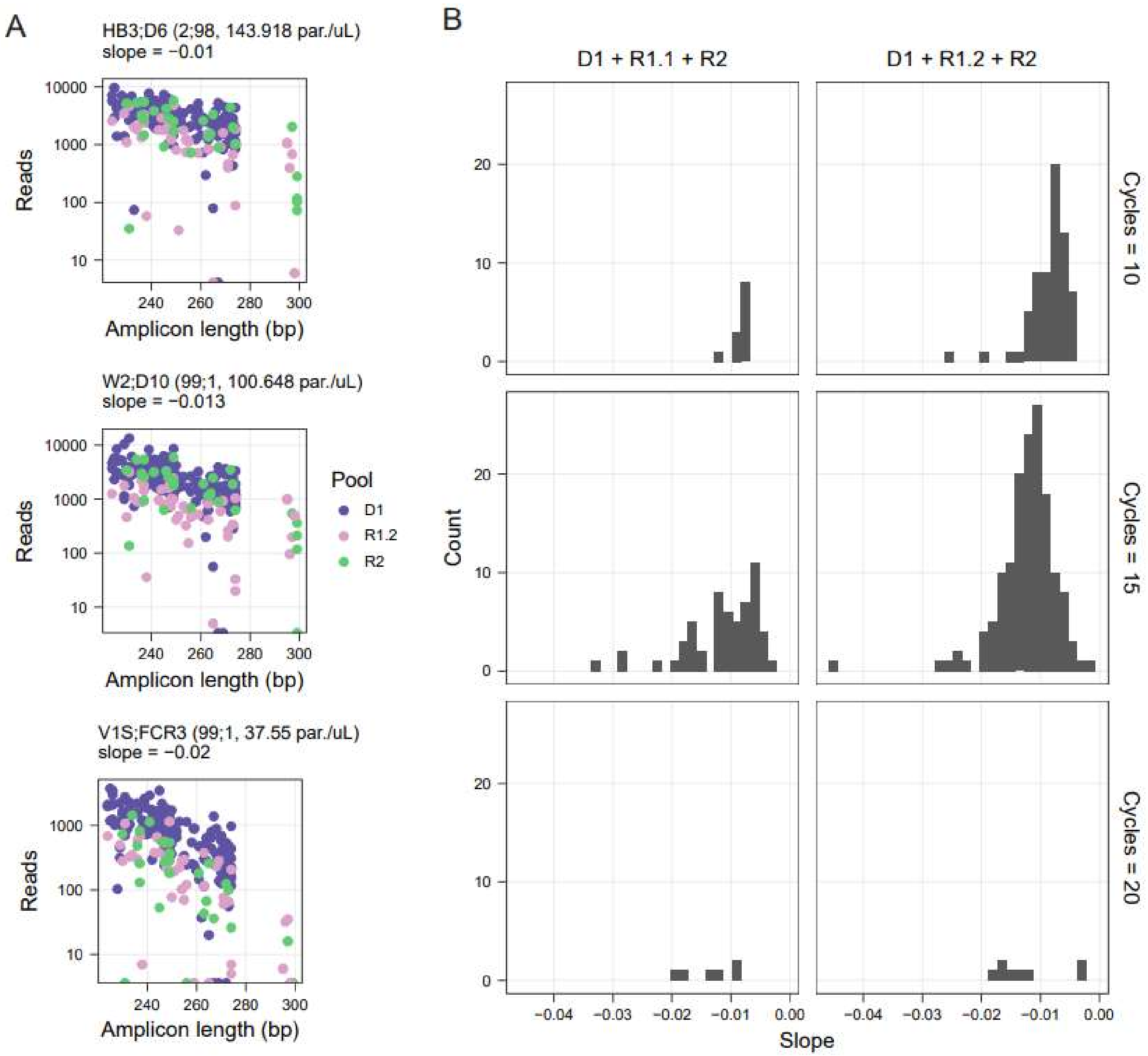


**Supplementary Figure 6. Depth of coverage is negatively correlated with amplicon length.**

**A.** Read counts for all targets in libraries made from mixed DBS controls with varying amplicon length amplification bias. The DBS control strains are indicated with its composition (e.g. W2;D10 2;98 is 2% W2, 98% D10) and parasite density. Note that libraries were prepared at the same time with 15 multiplex PCR cycles and the same reagent master mixes.

**B.** Distribution of slopes of the relationship between log10(read counts) and amplicon length in DBS controls libraries made in different laboratories and using different conditions shows they are predominantly negative and variable.

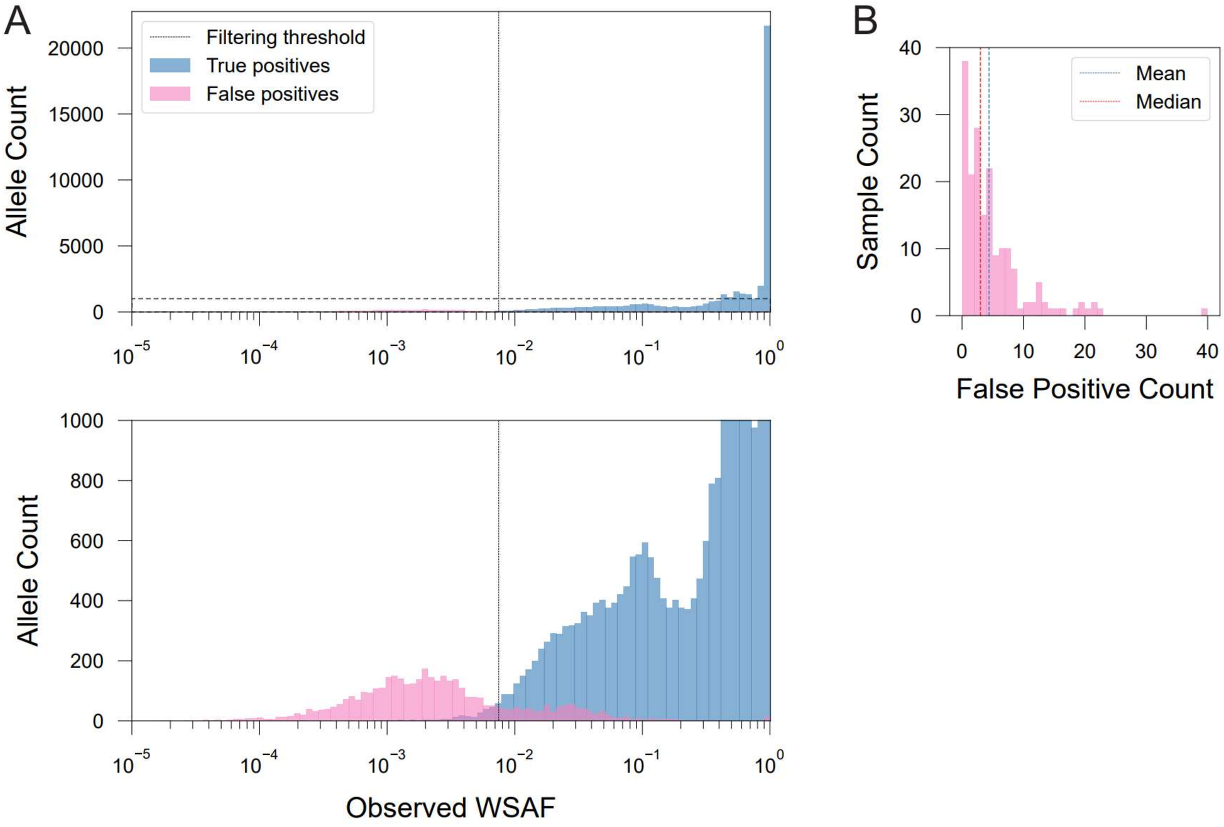


**Supplementary Figure 7**

**A.** Histogram depicting the distribution of observed alleles at varying WSAF (within locus and sample), with true positives (expected based on truth built from monoclonal controls) and false positives (not expected alleles) distinguished by different colors. The data was generated in five different laboratories from mixed DBS controls with known composition at varying strain proportions and parasite densities. The bottom panel is an inset of the top panel to better visualize the false positives. Based on these results, we implemented a filtering threshold of 0.75% within-sample allele frequency (WSAF) as it is approximately the point in which observed alleles become predominantly false.

**B.** Distribution of false positive allele counts per sample (mean 4.4, median 3 of a total of 161 targets). Alleles with ≤ 0.75% WSAF were excluded.

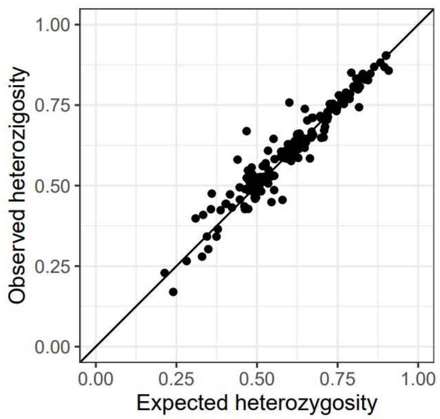


**Supplementary Figure 8.** Observed heterozygosity in samples from Mozambique is highly correlated with the expected heterozygosity. Expected heterozygosity was calculated from publicly available WGS data while the observed heterozygosity was obtained from a study that used MAD4HatTeR to generate allele data. Alleles used to generate the data in this figure are not masked in homopolymers and tandem repeats, resulting in higher heterozygosity values than the ones in Figure 4G.

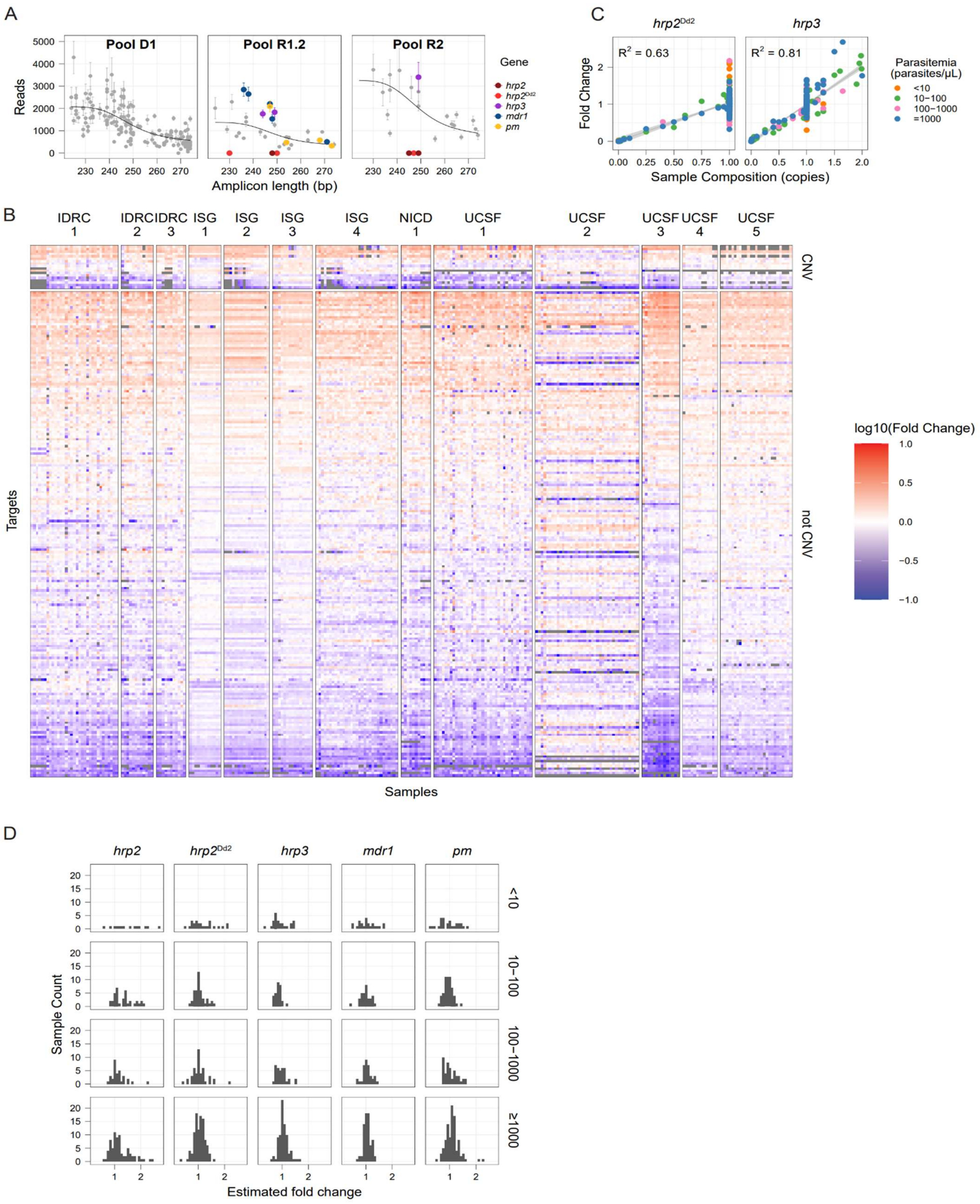

A
Pool D1
Pool R1.2
Pool R2
Reads
Amplicon length (bp)
Gene
hrp2
hrp2Dd2
hrp3
mdr1
pm
B
IDRC 1
IDRC 2
IDRC 3
ISG 1
ISG 2
ISG 3
ISG 4
NICD 1
UCSF 1
UCSF 2
UCSF 3
UCSF 4
UCSF 5
CNV
not CNV
Targets
Samples
log10(Fold Change)
C
hrp2Dd2
hrp3
R2 = 0.63
R2 = 0.81
Fold Change
Sample Composition (copies)
Parasitemia (parasites/μL)
<10
10−100
100−1000
=1000
D
hrp2
hrp2Dd2
hrp3
mdr1
pm
<10
10−100
100−1000
≥1000
Sample Count
Estimated fold change


216

**Supplementary Figure 9**

**A.** A generalized additive model is used to correct for biases in sequencing depth due to amplicon length and differences in primer pools. In addition to that, the raw reads for targets of interest (shown in this figure) are corrected by amplicon-specific bias using fold changes obtained from controls known not to have duplications or deletions processed in the same laboratory within the same timeframe. The corresponding corrected data is shown in Figure 5A. The data corresponds to technical replicates of Dd2 (a strain with *hrp2* deletion and *mdr1* duplication) with similar total reads were used to estimate fold changes in targets in and around *hrp2*, *hrp3*, *mdr1* and *plasmepsin2*/*3* (*pm*). A generalized additive model (black line) was applied to raw reads. Note that there are two groups of *hrp2* targets, those that are deleted in field samples (*hrp2*)[3] and those also deleted in Dd2 ($hrp2^{Dd2}$). Mean reads are shown (N = 3); error bars denote standard deviation.

**B**. Sample batches show different biases in amplification that are not explained by primer pool differences or amplicon length biases. The residuals for each target using a generalized additive model accounting for primer pools and amplicon length are shown for the samples used in Figure 5, grouped by sample preparation batch (samples prepared in the same laboratory within a time frame). CNV (duplications and deletions) targets of interest are also grouped separately. We note that some sample batches have very different amplification bias profiles (e.g. UCSF 2), possibly due to differences in reagent manufacturing lots. Despite those differences, use of appropriate controls allows to detect fold changes in targets of interest.

**C**. Estimated fold change for *hrp2* targets only deleted in Dd2 ($hrp2^{Dd2}$), as well as data including samples with *hrp3* duplications (containing strain FCR3), in laboratory controls containing 1 or more strains at known proportions, or in field samples from Ethiopia[3] with known CNV genotype. Fold changes are obtained using the targets highlighted in A. Linear regression and $R^2$ values were calculated with data with parasite density > 10 parasites/µL.

**D**. Distribution of fold changes in controls known not to have a duplication or deletion show that the estimations are noisier in samples of lower parasite density and suggest that the method can be used for targets for which we do not have enough validation data (i.e. *plasmepsin2/3* duplications, *pm*) .

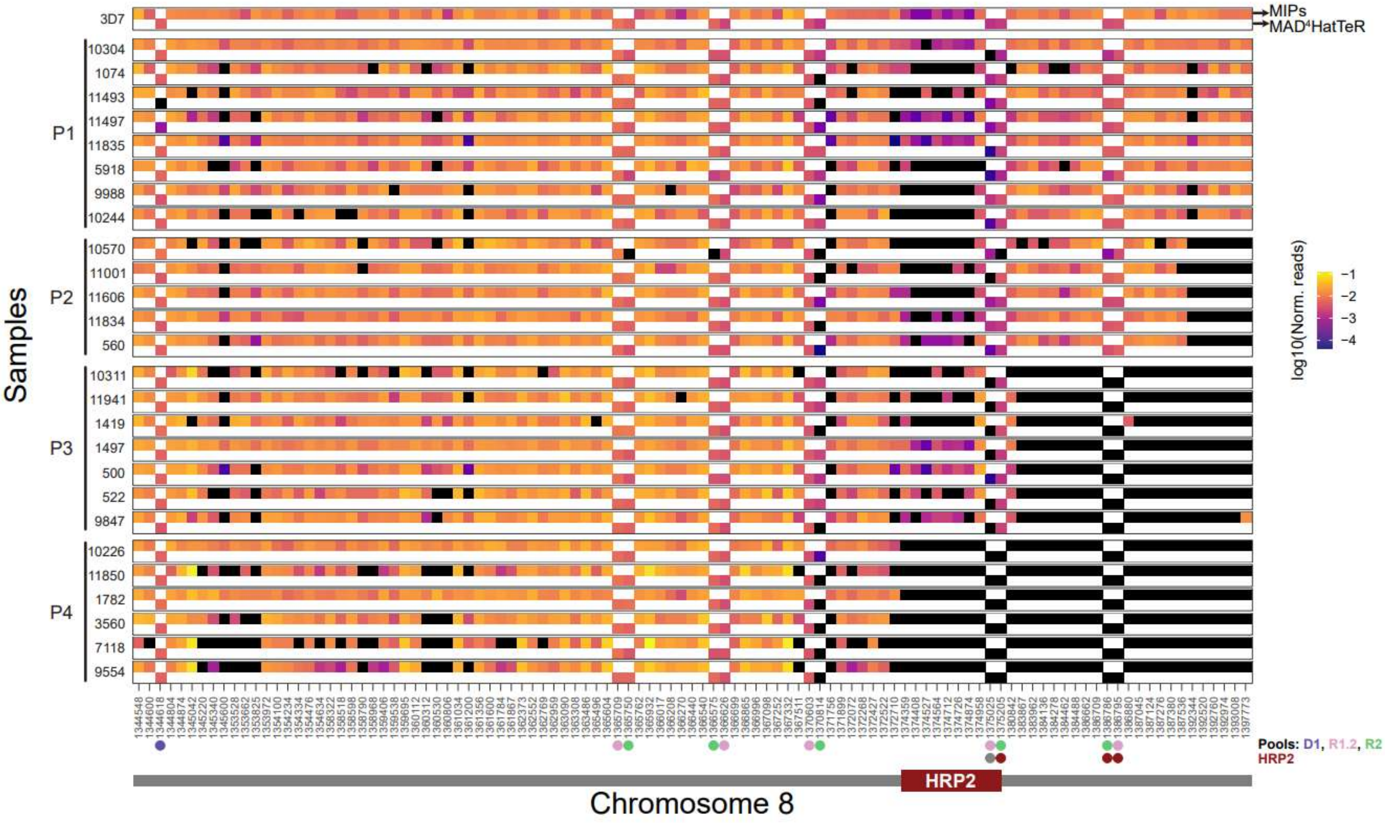
MIPs
MAD4HatTeR
3D7
P1
P2
P3
P4
Samples
log10(Norm. reads)
Pools: D1, R1.2, R2
HRP2
Chromosome 8

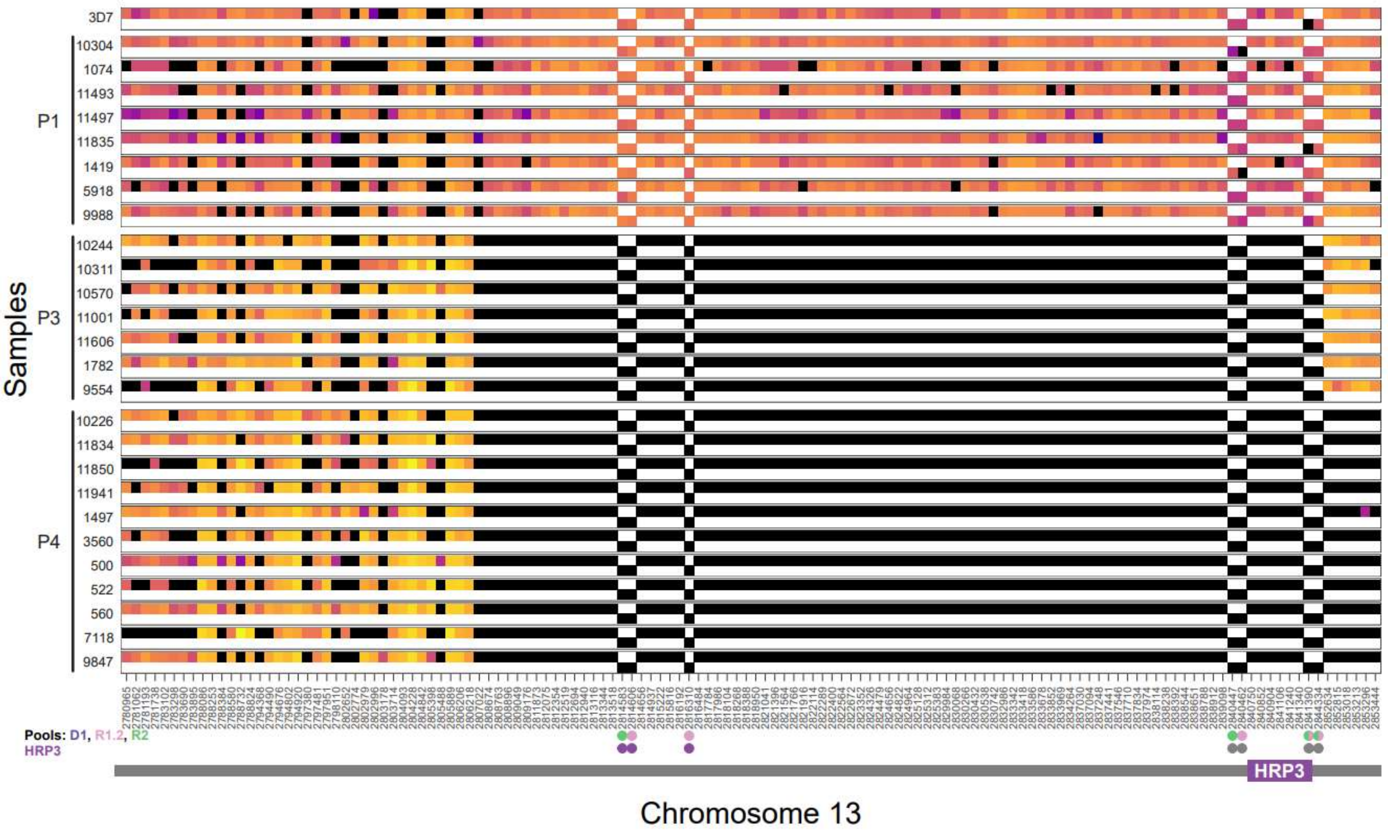
3D7
P1
P3
P4
Samples
Pools: D1, R1.2, R2
HRP3
Chromosome 13

**Supplementary Figure 10.** Comparison of data generated by molecular inversion probes (MIPs) and MAD$^4$HatTeR for field samples from Ethiopia previously shown to have *hrp2*/*3* deletions[3]. MIP data was generated in the original study, while MAD$^4$HatTeR data was generated from the same DNA extracts. Reads were normalized to the total number of reads in the chromosome (MIPs) or the primer pool (MAD$^4$HatTeR). Samples are grouped according to the breakpoint profiles (P1-4) as described by Feleke *et al*. Probe and amplicon positions are indicated by their start points. MAD$^4$HatTeR targets are indicated by a dot, with colors indicating the primer pool they belong to. Targets used in the generalized additive model to estimate fold changes are denoted by a dot in dark red or purple at the bottom of the plot. Targets that were excluded from the analysis due to variability are shown in gray. To estimate *hrp2* fold changes using MAD$^4$HatTeR data the model uses the maximum estimated fold change for either the target at the end of the *hrp2* gene (starting at position 1375205) or the two downstream targets combined. For *hrp3*, the estimated fold change is calculated on all three included targets. Note that chromosome positions are not to scale.

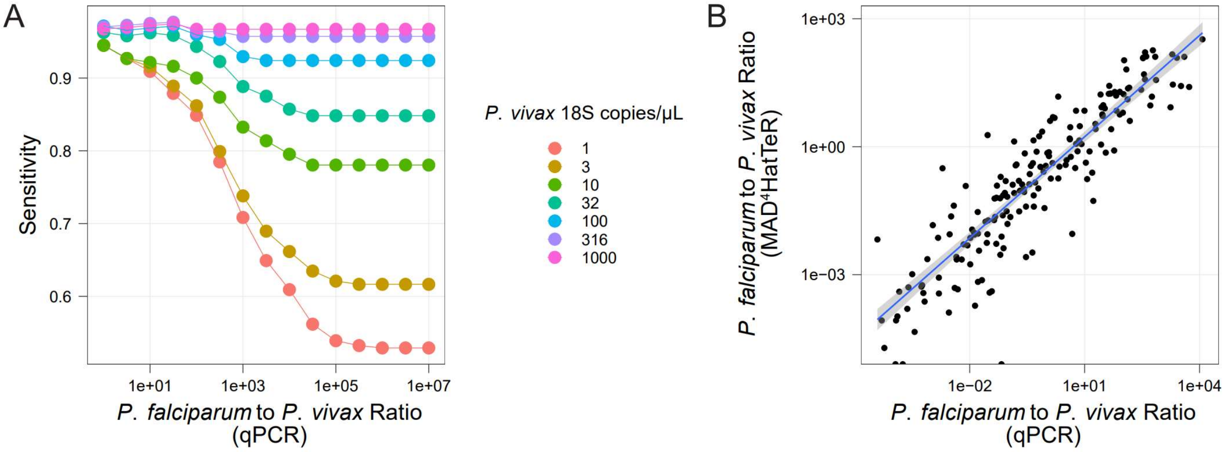


**Supplementary Figure 11.** MAD$^4$HatTeR's ability to detect *P. vivax* in *P. falciparum* coinfections.

**A.** Sensitivity of *P. vivax* detection by MAD$^4$HatTeR in 288 Ethiopian samples containing P. vivax and P. falciparum. 18S qPCR was used to confirm the presence of both species and to estimate the ratio of *P. falciparum* to *P. vivax* in each sample. Sensitivity was calculated excluding samples with a *P. falciparum* to *P. vivax* ratio greater than the values in the x axis and with *P. vivax* parasite density greater than the values indicated in colors.

**B.** The estimated *P. falciparum* to *P. vivax* using parasite densities (qPCR) and MAD$^4$HatTeR are correlated. Only samples with both species detected by both assays are included. $R^2 = 0.77$, $p = 2.2 \cdot 10^{-16}$.

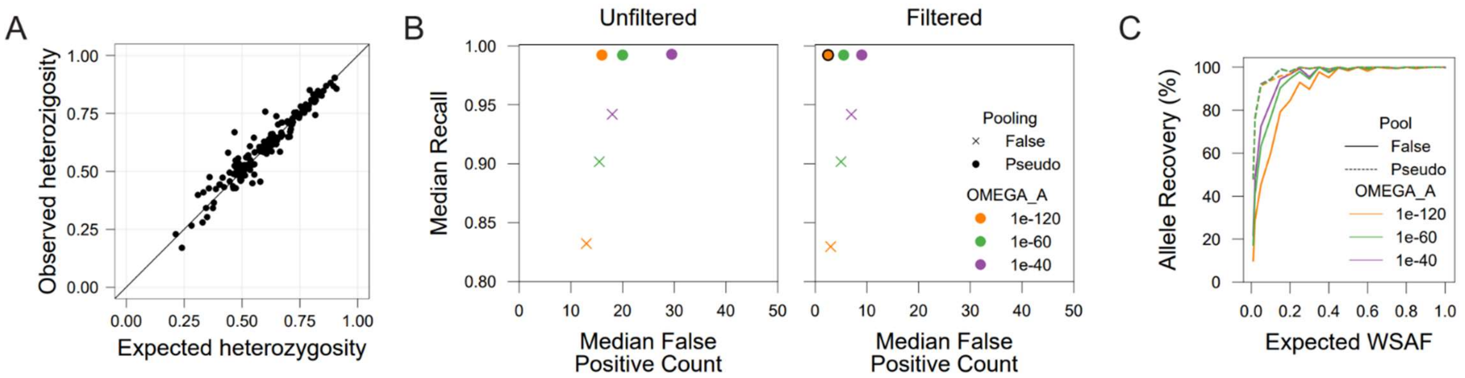


**Supplementary Figure 12. Bioinformatic pipeline benchmarking and optimization**

**A.** Homopolymer and tandem repeat masking increases precision but also removes real biological variation. Heterozygosity was calculated from publicly available WGS data for samples from Mozambique using raw alleles or alleles masked for homopolymers and tandem repeats. 14 targets showed a sizable decrease in heterozygosity.

**B.** DADA2, the denoising algorithm used in the bioinformatic pipeline, can be tuned to optimize sensitivity while minimizing false positives. The median proportion of expected alleles observed across samples and the median false positive count per sample was calculated for combinations of two DADA2 parameters. Decreasing OMEGA_A increases stringency, while using pseudo pooling allows for the retrieval of low abundance alleles observed in other samples in the sequencing run. Furthermore, post-DADA2 filtering of alleles based on their WSAF (using a 0.75% threshold, Supplementary Figure 7) decreases the median false positives. Chosen parameter settings used in this study, which maximize recall of expected alleles and minimize false positives, are OMEGA_A=1e-120, pseudo pooling and 0.75% filtering.

**C.** DADA2 parameters mostly affect the recovery of low abundance alleles, and the effect of OMEGA_A on sensitivity is minimized by pseudo pooling.

### Supplementary text

*Library preparation and sequencing*

The procedures are described according to the workflows at UCSF unless otherwise noted. Minor variations (e.g. equipment) were implemented at other institutions.

To extract DNA, we used the Chelex-Tween method as previously described[4]. Briefly, we punched 6 mm discs from DBS either into 1.5 mL microcentrifuge tubes or 2 mL round bottom 96 deep-well plates. We incubated the discs in 0.5% Tween 20 (Sigma Aldrich P9416) in 1X PBS (Corning 21-040-CV) overnight at 4 °C with constant shaking. We discarded the supernatant, washed the discs with chilled 1X PBS for at least 30 minutes at 4 °C with continuous shaking, and added 150 uL 10% Chelex 100 (Bio-Rad 1422822) in water. After incubating at 95 °C for 10 minutes, we centrifuged the samples for 5 min at 20,000 rcf and transferred the supernatant to 0.5 mL sample storage tubes. We stored extracted DNA at -20 °C.

To estimate *P. falciparum* parasite density, we ran an ultrasensitive qPCR varATS assay[5]. We made qPCR reactions with 1X Taqman Gene Expression, 0.8 µM forward (cccatacacaaccaaytgga) and reverse (ttcgcacatatctctatgtctatct) primers, and 0.4 µM varATS probe (6-FAM-trttccataaatggt-NFQ-MGB). We added 5 µL of extracted DNA for a total reaction volume of 25 µL. We ran qPCR reactions using the following settings: Pre-incubation: 2 minutes at 50°C; Initial Denaturation: 10 minutes at 95°C; Amplification: Denaturation: 15 seconds at 95°C, Annealing and Elongation: 1 minute at 55°C. We ran each reaction in the QuantStudio 3 (ThermoFischer A28567) for 60 reaction cycles and read fluorescence at the end of each cycle. In every plate, we ran standards at the following parasite concentrations in duplicates: 10,000, 1,000, 100, 10, 1, 0.1 and 0.05 parasites/µL. We estimated parasite densities using linear regression on standards using the QuantStudio Design and Analysis Software. Some participating laboratories (ISGlobal, CISM) performed an 18S qPCR[6] to estimate parasite density..

We ordered primers for multiplexed amplification already pooled at 5X concentration and used the CleanPlex Targeted Library Kit for 2-Pools Panels (Paragon Genomics). We mixed compatible modules with equivolume for multiplexed PCR.

We made multiplexed PCR reactions with 1X mPCR master mix and varied primer concentrations (0.125 - 1X). We used 6 µL of input DNA regardless of parasite density, and the total reaction volume was 10 µL. We ran mPCR reactions using the following settings: Initial denaturation: 10 min at 95 °C; Amplification: Denaturation: 15 s at 98 °C with 3°C/s ramping; Annealing and elongation: 5 min at 50°C with 2°C/s ramping. We repeated the Amplification steps for varying cycles (10-20). We combined mPCR products for the same sample using two incompatible primer pools and added 4 µL STOP buffer for a total volume of 24 µL. Alternatively, we added 10 µL TE buffer and 2 µL Stop Buffer for single-tube reactions. We purified amplicons (225-300 bp) and removed primer dimers using 1.3X CleanMag Magnetic Beads solution (Paragon Genomics). After incubating for 5 mins at room temperature, we washed magnetic beads twice with 70% ethanol in water and eluted DNA in 10 µL TE buffer after ethanol evaporation. We degraded non-specific DNA products with Digestion Reagent for 10 minutes at 37 °C and purified amplicons with 1.3X CleanMag Magnetic Bead as above after stopping the reaction with 2 µL Stop Buffer. We further amplified eluted DNA with 1X 2$^{nd}$ PCR Master Mix and 0.5 µM indexing primers. Indexing primers were either custom-made (comprising the i7 or i5 sequences and TruSeq adapters flanking 12 bp indexes) or CleanPlex Plated Unique Dual-Indexed PCR Primers for Illumina (Paragon Genomics). We ran indexing PCR reactions using the following settings: Initial denaturation: 10 min at 95 °C; Amplification: Denaturation: 15 s at 98 °C with 3°C/s ramping; Annealing and elongation: 1:15 min:s at 50°C with 2°C/s ramping. We repeated the Amplification steps for 15 cycles.
After initial optimization, we identified the following optimal conditions for amplification: (1) 15 cycles and 0.25 X primer concentration for mPCR for parasite densities equivalent to ≥ 100 parasites/µL in DBS; (2) 20 cycles and 0.125 X primer concentration for mPCR for <100 parasites/µL. We note that while 20 cycles do not significantly increase library yields, anecdotal information across laboratories indicates that 20 cycles with lower primer concentrations increases the success rate of amplification from samples with low parasite densities.

After amplification, we pooled samples accounting for expected differences in yield. We pooled DBS-control samples based on starting parasite density with the following schema: 3 µL for 10,000 parasites/µL; 6 µL for 1,000; 12 µL for 100; 20 µL for 10; and 30 µL for 1. We pooled up to 96 samples with 10 µL for each sample where the initial parasite density was unknown. We purified each pool to remove primer dimers further using 1X CleanMag Magnetic Bead Solution as above and eluted in 40 µL TE buffer. If necessary, to completely remove primer dimers, we gel-purified each pool in a 2.5% agarose gel with SyberSafe 1X (S33102) alongside a 100 bp ladder (GeneRuler SM0242). We ran the gel for 60 minutes at a constant voltage set to 140 V. We excised the 400 bp band and purified DNA using the NEB Monarch Gel Extraction Kit (New England Biolabs, T1020L). We performed a second gel purification if the sample had more than 5% integrated area in the 150-250 bp region.

We evaluated library preparation performance with capillary electrophoresis using an Agilent 4150 TapeStation System using D1000 Reagents (5067-5583) and ScreenTape (5067-5582). We recommend using these to evaluate library yield.

We ran pools with 5 or 10% PhiX and observed no large differences with either concentration.

*A bioinformatic pipeline for MAD$^4$HatTeR*

Inputs to the pipeline are (a) already demultiplexed sample-based fastq files; (b) a file containing information of each amplicon, including primer sequences and genomic locations; (c) a list of resistance markers; (d) a reference genome. The pipeline consists of multiple processes:

(1) CREATE_REFERENCE_SEQUENCES: Reference sequences are used to identify polymorphisms in the amplicon data and generate pseudo-CIGAR strings. This module creates reference sequences for each amplicon using a user-provided genome and coordinates supplied in the amplicon information table and stores reference sequences in a fasta file. This module is skipped if the user provides reference sequences for each amplicon.

(2) MASK_SEQUENCES: (optional) Regions in the reference sequences that contain short tandem repeats, identified using Tandem repeats finder[7], or homopolymers longer than a user-determined threshold (default is 5), are masked. These low-complexity regions are known error-prone sites in PCR and Illumina sequencing[8]. Masked sequences have each nucleotide base replaced with an ‘N’ character.

(3) CUTADAPT: Fastq files are filtered based on read quality, and primer dimers are removed based on the presence of Illumina adapter sequences in the read using cutadapt. Here, we use the following parameters: `-e 0` - This sets the allowed number of mismatched bases to zero when identifying adapters in the sequence.; ’--no-indels` - This prevents adapters from being identified if there are any insertions or indels in the sequence; `--minimum-length` - This is set to 100, and removes reads that are shorter than this threshold; `--untrimmed-output` / `--untrimmed-paired-output` - Any forward

reads that do not contain the Illumina adapter in their sequence are written to file. Adapter dimers are also captured for quality control.

These reads are further demultiplexed to generate individual fastq files for each amplicon in each sample using a second cutadapt step to identify primer sequences flanking the reads. We use the same parameters above to identify primer sequences (only exact matches anchored to the beginning of the read are allowed), and filter reads that are less than 100 bases after primer removal. Error filtering is conducted based on sequencer: if the data is generated with a MiSeq, `--trim-n -q 10` is set to remove ambiguous base calls and remove low-quality bases (less than 10) on the 3' end of the read. If the data is generated with a NextSeq, '--nextseq-trim=20' is used to remove low-quality bases (less than 20) and high-quality G bases found on the 3' end of the read.

(4) QUALITY_REPORT: Per-sample and per-amplicon summaries are made based on three numbers: reads inputted, reads that passed filters in the first cutadapt step, and reads that were demultiplexed using primer sequences.

(5) DADA2: DADA2's filterAndTrim, learnErrors, dada, mergePairs, and removeBimeraDenovo functions are used to filter fastqs demultiplexed by amplicon and sample, parameterize an error model based on read quality, infer 'true' variants (amplicon sequence variants, ASV) and remove chimeras in each fastq file. During filtration, we remove reads that have more than at least two expected errors (MAXEE=2) to generate consistent error models between dataset runs. Forward and reverse reads with a PHRED quality score of less than 5 are also truncated (trunqQ=5), and all reads have their leftmost base trimmed (trimLeft = 1). All reads less than 75 bases (after truncation) are filtered out, and the rest of the filtration parameters are set to the package defaults. To train the error model, samples are randomly selected until the default minimum number of bases requirement is met (nbases=1e8), and all other parameters use the package defaults. Finally, the allele inference algorithm uses custom `OMEGA_A=1e-120` and `POOL=pseudo` settings to identify real biological sequences from the dataset and was run in SELF_CONSIST mode to maintain error model estimation after each sample composition inference step.

Merging of clustered forward and reverse reads is done with custom code that identifies targets without a minimum overlap and uses DADA2's mergePairs function. Targets with an overlap of at least 10 base pairs are merged; reads are concatenated otherwise.

(6) DADA2_POSTPROCESSING: DADA2 ASV data is passed onto custom code to rearrange the data and filter out ASVs by alignment to reference sequences using

Needleman-Wunsh global alignments with the Biostrings package from Bioconductor. Alignment scores are calculated for each comparison and the best alignment is kept, penalizing mismatches (-1) and gaps (openings = -8, extension = -5). A default alignment score of 60 is used to filter out particularly poor alignments caused by off-target sequences and can be adjusted by the user as needed. Optionally, homopolymers and tandem repeats can be masked to reduce spurious output from sequencing errors. Masked sequences can be provided by the user, or can be created using Tandem repeats finder[7] as part of the pipeline, which will identify short tandem repeats using Smith-Waterman local alignments and is controllable by user settings (defaults: period_size = 3, alignment_score = 25). Homopolymers can also be masked depending on the set user threshold (default: homopolymer_threshold = 5), and will mask leading and trailing bases to hide substitution errors caused by phasing. While masking is optional, benchmarking suggests it can improve precision and sensitivity results. To store alignment and masking information, we used a pseudoCIGAR representation of the aligned ASVs to the masked references. This pseudoCIGAR indicates:

(a) Masked regions: the start of the masked region followed by its length (α+βN, where α is the start position and β is the length)
(b) Substitutions: the substitution position and the base it's substituted to (αX, where α is the start position and X is the substituting base)
(c) Insertions: the start of the insertion and the sequence inserted (αI=X`, where α is the start position, I indicates an insertion, and X` is the base or bases inserted)
(d) Deletions: the start of the deletion and the sequence deleted (αD=X`, where α is the start position, D indicates a deletion, and X` is the base or bases deleted)
(e) Perfect match. Denoted by the period symbol (.).

*Bioinformatic pipeline benchmarking*

We benchmarked the bioinformatic pipeline to identify optimal parameters for sensitivity and precision using data generated from DBSs containing combinations of 9 cultured strains (FCR3, V1S, W2, U659, D10, D6, HB3, Dd2, 3D7). To build a set of expected (true) alleles (Supplementary Tables 9 and 10), we used at least 1 monoclonal sample for each strain, using the pipeline with restrictive DADA2 parameters (OMEGA_A=$10^{-120}$ and pool=false). After filtering out pseudoCIGARs with < 10 reads, the true allele for each strain at each locus was identified as the major (>90%) pseudo cigar at that locus.

DADA2 parameters OMEGA_A and pooling can have a large impact on both sensitivity and precision. OMEGA_A sets the threshold for DADA2 to classify an entity as abundant enough to form its own ASV. Setting this threshold too conservatively can reduce the number of false positives, but it also decreases sensitivity, potentially overlooking rare variants. On the other hand, sample pooling strategies (none, pseudo-pooling, or pooling) can help inform the inference of ASVs in each sample with the ASVs observed in others. We discarded true pooling as it is highly computationally intensive. When pseudo pooling is enabled, DADA2 performs a second round of inference, using ASVs identified in the first round across all samples within the run as a prior. This iterative approach can enhance the sensitivity of ASV detection for minor alleles that are common within a run (and ultimately can be defined for a population) but it can also lead to the retrieval of spurious sequences or, more importantly, low-level contaminants. In addition to these DADA2 parameters, we built a masking algorithm into the pipeline to identify tandem repeats and homopolymers and ignore any variation observed within them. This strategy removes true biological variation (14 targets showed a > 0.1 drop in heterozygosity when masked, Supplementary Figure 12A). Nevertheless, it allows us to discard false positives in hotspots; we observed a drop in false positives per sample from 28 to 21. Further characterization of real biological variation and sequencing and PCR artifacts, will be needed to fine-tune masking parameters, including masking regions *ad hoc*. The pipeline also reports unmasked ASVs to give the users the option to use either. We proceeded with masking for the rest of the analysis.

To fine-tune the balance between stringency and rare variant recovery afforded by DADA2 we processed sequencing runs with combinations of OMEGA_A ($10^{-40}$,$10^{-60}$,$10^{-120}$) and pooling methods (pseudo, false). To evaluate its impact, we estimated precision and sensitivity using targets in the diversity module D1 genotyped in 158 controls (72 unique combinations of strains at different proportions) sequenced across 12 sequencing runs in the 5 participating sites. Most runs included field samples prepared with MAD$^4$HatTeR that are not part of this study which could be sources of contamination and lead to false positives.  Sensitivity was defined as the median(TP/(TP+FN)) across all loci in a sample; the median sensitivity across samples was then used to evaluate the effect of different parameters. Precision was evaluated with the median number of false positives per sample. The truth generated from the monoclonal samples was used to classify pseudoCIGARs returned by the pipeline into true or false positives. False negatives were identified if the true pseudoCIGAR was not observed, irrespective of the presence of other true or false positive pseudoCIGARs in that locus and sample. We noticed that irrespective of parameters, a large amount of very low abundance false positives were retrieved, most of which were not expected as the minimum minor WSAF in the set was 1%. Thus, we also imposed different within-sample allele relative cutoffs to filter out alleles. A WSAF of 0.75% minimized false positives while maintaining false negatives low (Supplementary Figure 12B). More restrictive OMEGA_A values decreased sensitivity, but this effect was counteracted by pseudo-pooling; in the presence of pseudo-pooling, OMEGA_A had little to no effect on sensitivity, while it still improved precision (Supplementary Figure 12B). Pseudo-pooling improved the recall of alleles expected below 40%, leading to an overall improvement in sensitivity, while marginally increasing the number of false positives (Supplementary Figure 12C).

We note that the interplay between these parameters and the experimental processes can lead to biased results. For example, the use of pseudo-pooling can disproportionately increase the retrieval of contaminants because inference is informed by the presence of ASVs in other samples within the same run. Indeed, the increase of false positives was more marked in samples that already had higher false positive rates, suggesting that these are generally contaminated at low levels. Using absolute and relative abundance filters minimizes this effect.

*Copy Number Variation (duplications and deletions)*

We used the following laboratory strains to benchmark CNV detection using MAD$^4$HatTeR data: *hrp2* deletions in Dd2 and D10, *mdr1* duplications in Dd2 and FCR3, *hrp3* deletion in HB3, and *hrp3* duplication in FCR3[9]. We also used a set of field samples previously shown to have deletions in and around *hrp2* and *hrp3*, including multiple breakpoints[3]. For sensitivity analysis using field samples, we estimated COI using moire[10] and excluded polyclonal samples due to the uncertainty in their true genotypes. Two field samples were excluded from the analysis due to discordance in breakpoint classification, possibly due to sample mislabeling and sequencing depth, respectively.
We estimated read depth fold changes from data for each gene of interest (*hrp2*, *hrp3* and *mdr1*). We did not have sufficient data to validate CNVs in plasmepsin 2 and 3. We excluded targets with high variability in depth across samples (N = 220, Supplementary Table 6). Only samples with at least 180/220 targets with > 10 reads each, and with < 100-fold difference in

depth between reactions were used. Filters left the following number of targets for each gene: 6 for *hrp2*, 3 for *hrp3*, and 5 for *mdr1*. Samples with deletions around *hrp2* can be clustered based on their breakpoint[3] (Supplementary Figure 10). One target (Pf3D7_08_v3-1375205-1375450-2) used in the analysis allows the detection of deletions in *hrp2*, while 2 other targets can detect deletions downstream of *hrp2*. That downstream deletion is observed in all *hrp2* deletions[3]. To increase specificity and sensitivity and ensure true deletions in the *hrp2* gene were detected, we used the maximum fold change observed for either Pf3D7_08_v3-1375205-1375450-2 or the downstream targets. The other three *hrp2* targets are in an upstream region deleted only in Dd2 and they are grouped independently.

Fold changes were estimated using a generalized additive model on reads (dependent variable) per target and sample. The model incorporated a smooth spline function of amplicon length (due to the correlation between depth and length, Figure 5A) with basis dimension $k$ set to 4, and a categorical variable for each primer pool and linear terms for predictors that identify the targets corresponding to each gene of interest (*hrp2*, *hrp2* Dd2 targets, *hrp3*, and *mdr1*). Biases in amplification that were not explained by length or primer pools varied across sample preparation batches. Thus, we first calculated the median residuals for the targets in each gene in controls within a batch (defined as data generated in the same laboratory within a time frame that did not lead to stark changes in biases within the controls) and corrected the reads for those targets in each of the queried samples. Controls were monoclonal 3D7 where available, or other strains known to lack CNV in the assayed genes. For one batch of samples a mixed control with 98% non-CNV was used as a 100% non-CNV control was not available. Corrected reads in queried samples were then applied to the model as explained above and the coefficient for each group of targets was reported as a proxy for the gene depth fold change.
